# Nation-scale recovery of complete environmental plasmids pushes the boundaries of plasmid diversity through long-read sequencing

**DOI:** 10.64898/2026.10.06.756670

**Authors:** Bertil Sørensen, Connor L. Brown, Mantas Sereika, Søren Heidenbach, Philip Hugenholtz, Mads Albertsen, Francesco Delogu

**Author notes:** Shared first authorship (Bertil Sørensen, Connor L. Brown).

## Abstract

Plasmids are important agents of prokaryotic evolution and yet existing reference databases remain biased towards cultivable species, limiting our knowledge of their function and ecology in the environment. Here we present a major expansion of known plasmid diversity, through curation and annotation of 8,744 topologically complete plasmids from 9,972 short- and 154 long-read metagenomes as part of the *Microflora Danica* project. These were found to increase plasmid taxonomic unit diversity by 15% and 99% relative to all and environmental plasmids, respectively, from public sources. In particular, we identify a novel lineage of plasmids in the genus *Methanosarcina*, encoding genes associated with methanogenesis and cell surface properties, expanding our understanding of chromosome-plasmid complementarity and its potential contribution to host biology. Combining our plasmids with public sources, we curated a set of 17 replication-associated proteins that could be found in 82% of known plasmids and can serve as hallmarks for future detection of plasmids. We present this resource, and a tool for confirming topological completeness of plasmid assemblies, as a toolkit for future plasmid discovery.

## Introduction

Plasmids are partially autonomous DNA molecules that replicate as extrachromosomal elements within their host cells. Their replication is often governed by regulatory mechanisms distinct from chromosomal replication yet typically depends on the host’s replication machinery^1^. Plasmids are key drivers of antimicrobial resistance (AMR) through their role in horizontal gene transfer (HGT) permitting non-parental transmission of AMR genes between microbial lineages. Because of their importance to human health, plasmids associated with AMR are among the most thoroughly studied mobile genetic elements (MGEs)^2,3^. Beyond AMR, however, plasmids are increasingly recognized as vectors of broader ecological function, facilitating the transfer of genes involved in nutrient cycling, stress tolerance, and metabolic diversification^1^. These traits are important to the ecology of diverse environmental microbiomes such as soil, where the ecological functions encoded by the MGE pool remain largely unexplored^4^.

Plasmids are frequent hotspots for recombination and subject to complex copy number dynamics, therefore often yielding to highly fragmented assemblies from conventional short-read assemblers^5^. Long-read sequencing can mitigate this fragmentation, spanning repetitive and variable regions, leading to complete plasmid sequences that short-read assembly alone could not resolve^6^. However, long-read assembly introduces its own artefacts, including premature circularisation that results from algorithms misidentifying overlapping ends of a linear sequence due to repeats or structural variations^7,8^. Detecting plasmid-derived contigs from fragmented assemblies is also challenging^5^. Tools for identifying plasmid sequences often leverage sequence features (e.g., *k*-mers), hallmark genes, or a combination of the two^9^. For example, geNomad uses a hybrid strategy that combines *k-*mers with protein families found to distinguish plasmid, viral, and chromosomal sequences. However, frequent gene exchange and similarities to other elements (e.g., integrative or transposable elements) blurs boundaries between categories^10^. All classification approaches are further constrained by the plasmids in their training data. Major repositories such as PLSDB and IMG/PR remain heavily biased towards clinically relevant, human-associated plasmids^11,12^. PLSDB is compiled primarily from cultured isolates and IMG/PR from genomes and metagenomes, yet both are dominated by human-associated sources, leaving plasmids from natural environments substantially underrepresented. This margin of missing diversity was already noted a decade ago but remains a fundamental barrier to our knowledge of plasmid diversity^13,14^.

Classifying plasmids is fundamental to tracking how they (and their traits) spread through microbial communities. Efforts to classify and compare plasmids have evolved considerably alongside sequencing capacity^15^. Early groupings were based on differences in replication machinery (replicon typing) and transferability due to plasmid incompatibility (incompatibility groups)^9,16^. With increasing sequencing data, the presence of distinct replication-associated proteins (RPs) have come to serve as markers for these groupings. However, while these classification schemes remain important for plasmid nomenclature, the principles underlying them have proved difficult to translate to the diversity of plasmids revealed by sequencing^15^. More recent frameworks such as plasmid taxonomic units (PTUs) leverage average nucleotide identity (ANI) to facilitate large-scale comparative genomics of plasmids, revealing a chaotic landscape of mosaic replicon architectures and a substantial fraction of putative plasmids lacking detectable replication-initiating RPs^17,18^. Nishimura *et al*.^18^ recently addressed this in part by demonstrating shared evolutionary ancestry among plasmids encoding winged helix-turn-helix (WH) RPs. This single RP superfamily was associated with 57% of plasmids in PLSDB and 5,369 plasmids with no annotation from standard replicon-typing tools. Yet, this leaves a significant fraction of plasmid diversity without classification, particularly those encoded by archaea and other chiefly environmental bacterial lineages that do not use a WH replication initiator. RPs are attractive markers for identifying plasmids as they serve an essential role in plasmid replication and thus define the most basic plasmid function^1^. Thus, they may serve as strong markers of plasmid replication mechanism, host, and the macroevolutionary relationships of plasmid lineages.

Among the least represented plasmids in public databases are those of archaea^11^, despite their central role in major biogeochemical processes such as methanogenesis. Methanogens such as *Methanosarcina acetivorans* represent the main biological source of methane, encompassing 70% of global methane emissions^19^. Methanogens have been consequential across Earth’s history. Rothman *et al*.^20^ proposed that the horizontal acquisition of genes enabling acetoclastic methanogenesis in *Methanosarcina* supported a major expansion of methane production, contributing to carbon-cycle disruption and the end-Permian mass extinction 252 million years ago^20,21^. Such acquisitions may not be exceptional in this genus, as roughly a quarter of the *M. acetivorans* genome carries signatures of horizontal gene transfer, implying an extensive mobilome with the potential to expand or support host metabolism and ecology^22^. Despite these findings, little is known about *Methanosarcina* plasmids, with only two described in the literature^23,24^ and five others deposited in GenBank [accessions CP009527.1, CP009529.1, CP009525.1, CP009523.1 and CP009519.1]. Their roles in host biology remain uncharacterised.

Here, we used sequencing data from the Microflora Danica (MFD) project, comprising 9,972 short-read and 154 long-read metagenomes across Danish soil, sediment, and water environments, to recover a national-scale catalogue of environmental plasmids. We recovered and curated a high-quality set of environmentally-derived plasmids by assessing topological completeness through read-mapping and requiring plasmid hallmark genes, and placed them in their taxonomic and evolutionary context relative to publicly available plasmids. The resulting catalogue represents a major expansion of plasmid diversity, with most of the recovered plasmids representing unique and novel replicons. This was demonstrated by sequence-based clustering into plasmid taxonomic units (PTUs) and distant PTUs (dPTUs), whose biological coherence was assessed by examining conservation of backbone-associated and accessory traits within these groups. We further curated a set of replication-associated proteins from the combined plasmid set to aid plasmid identification, and characterise a novel archaeal plasmid lineage associated with *Methanosarcina* that encodes genes putatively associated with host methanogenesis and ecological adaptation.

## Results

### Recovery and curation of a diverse Danish soil plasmidome

To expand our understanding of plasmid diversity in natural environments, we identified complete, high quality plasmids from the Microflora Danica (MFD) dataset. Assembled contigs were screened using four published plasmid prediction tools (geNomad, Plasmer, viralVerify, PLASMe)^25–28^, yielding 6,388,240 and 1,561,754 putative plasmidial contigs from Illumina and ONT assemblies, respectively (**Fig. 1**).

**Figure 1:**
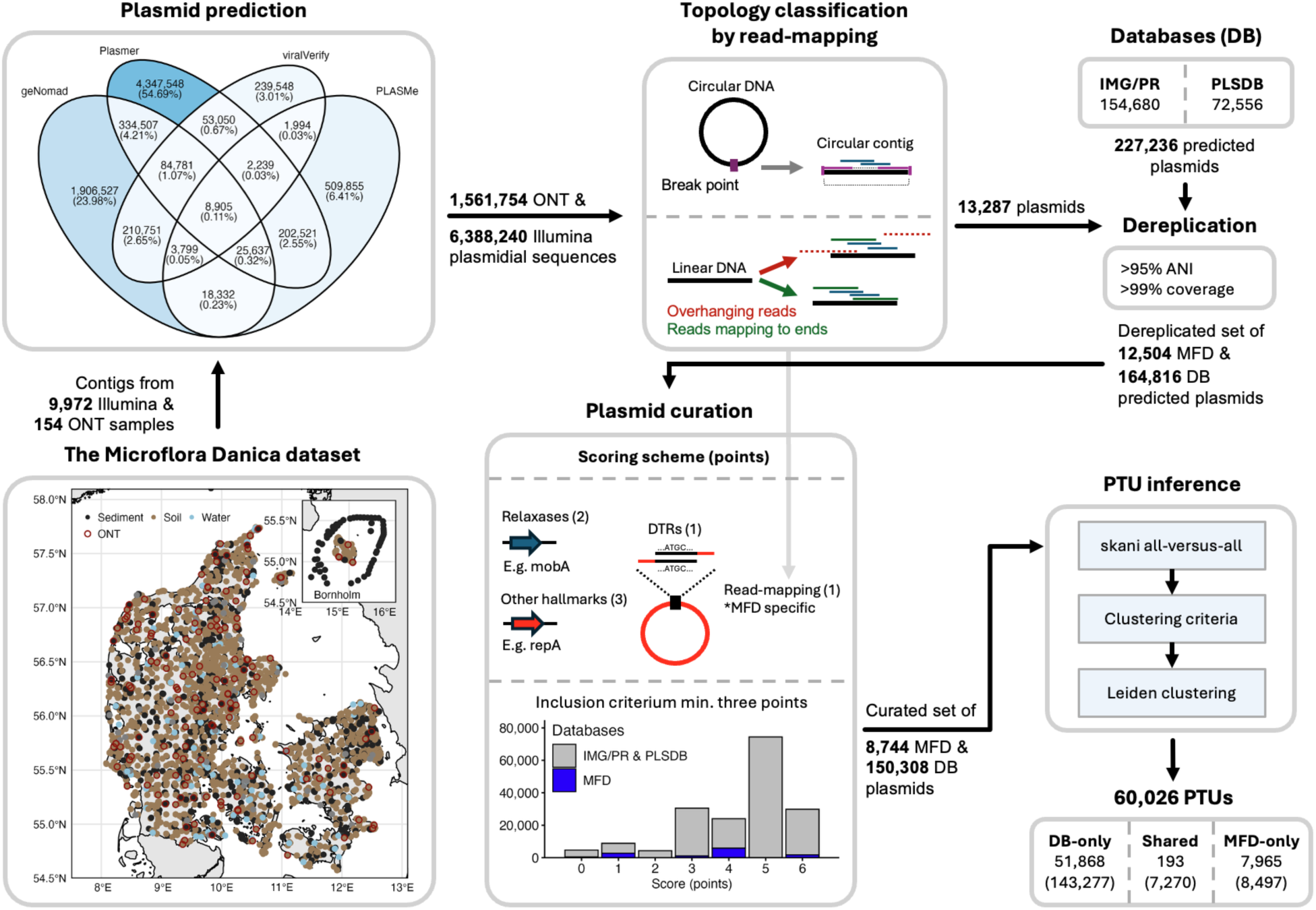
Contig assemblies from the Microflora Danica (MFD) dataset (9,972 Illumina and 154 Oxford Nanopore Technologies (ONT) metagenomes) were used as input for plasmid prediction. Contigs predicted as plasmid by at least one tool were then screened for topological completeness (circular or linear) by mapping the reads back to the assemblies. Plasmids classified as complete were dereplicated together with the reference collection of putatively complete plasmids from IMG/PR and PLSDB. The resulting plasmids were curated to retain a high-quality set of plasmids that were at minimum topologically complete and encoded a plasmid hallmark gene. In the scoring scheme, points were assigned for relaxases (+2), other plasmid hallmark genes (+3), direct terminal repeats (DTRs; +1) and read-mapping based completeness (+1; for MFD plasmids only). Plasmids scoring at least three points were retained. Sequence characteristics of the curated plasmid set are summarized in **Supplementary** Fig. 1.

There was remarkable disagreement among the plasmids predicted by the different tools, with only 8,905 contigs (0.11% of the total set) predicted by all tools **(Fig. 1)**. Of the four tools, Plasmer predicted the highest number of plasmidial contigs (comprising 4,347,548 plasmids or 54.7% of all predictions) followed by geNomad (1,906,527 plasmids or 24% of the total set). Of 55,029 contigs reported as circular by metaFlye^29^, 13,287 were supported as topologically complete by read-mapping, reflecting the inherent difficulty of resolving circular topology in complex metagenomes and underscoring the value of independently assessing assembler-reported circularity^8^. To focus our analysis on high quality plasmids, we devised a scoring scheme (inspired by IMG/PR^12^). Scores were assigned based on the presence of plasmid-specific genes (e.g., *mobA*, *repA*) and evidence of completeness, provided either by read-mapping (for MFD plasmids) or the presence of direct terminal repeats (DTRs). The scoring system required a minimum of three points per plasmid for inclusion **(Methods)**. A total of 8,744 dereplicated MFD plasmids (75.6% of which were recovered from ONT assemblies) and 150,308 dereplicated reference (IMG/PR + PLSDB) plasmids were retained after scoring and dereplication, henceforth referred to as “DB”. The combined and dereplicated MFD and DB dataset clustered into 60,026 PTUs. Additional sequence characteristics are shown in **Supplementary Fig. 1**.

Nearly all MFD plasmids belong to MFD-only PTUs (97% out of 8,744), indicating that most of our plasmids were not represented in the reference datasets **(Fig. 1).** Furthermore, among the MFD plasmids, 7,622 (88% of 8,744) formed singletons, compared to 37,207 of 150,308 database plasmids (25%). Thus, at the PTU-level, the majority of MFD plasmids have not been previously observed and are mostly unique relative to one another **(Supplementary Fig. 2).** Overall, this translated to an expansion of PTU-diversity by 15% at the PTU-level relative to the reference collection and 99.9% when only considering plasmids from environmental samples (i.e., IMG/PR plasmids with environmental ecosystem metadata flags). We also compared our plasmids with putatively complete plasmids from a recent global soil study^4^. Only three PTUs contained plasmids from both studies, and all three also contained plasmids from the reference collection **(Supplementary Fig. 3)**. Together, these results highlight a high degree of novelty and uniqueness among soil plasmids and indicate that current reference collections capture only a fraction of the underlying soil plasmid diversity.

## Danish terrestrial plasmids span diverse taxonomic units and replicon families

Few MFD plasmids could be clustered among DB PTUs (70% ANI and 50% AF). Thus, we attempted to identify more distant relationships between plasmid families by using average amino acid identity (AAI) and percent of conserved proteins (POCPs) to create distant (d)PTUs (70% AAI and 50% POCP). These measures have been used as an alternative to ANI among more distantly related genomes particularly where alignment coverage is insufficient to facilitate comparisons^30,31^. However, while utilizing dPTUs increased the number of MFD plasmids that could be grouped with DB plasmids by about ten-fold (2,282 vs. 247 at the PTU-level) (**Supplementary Fig. 4**), there remained a high degree (73%) of novelty even at this broader grouping **(Fig. 2A)**. This novelty also extended to protein sequence diversity, with MFD-only plasmids contributing 106,568 protein clusters absent from the reference collection at 30% amino acid identity **(Fig. 5B)**. Notably, this diversity persisted across broader PTU and dPTU groupings, while the reference collection became increasingly consolidated.

**Figure 2.**
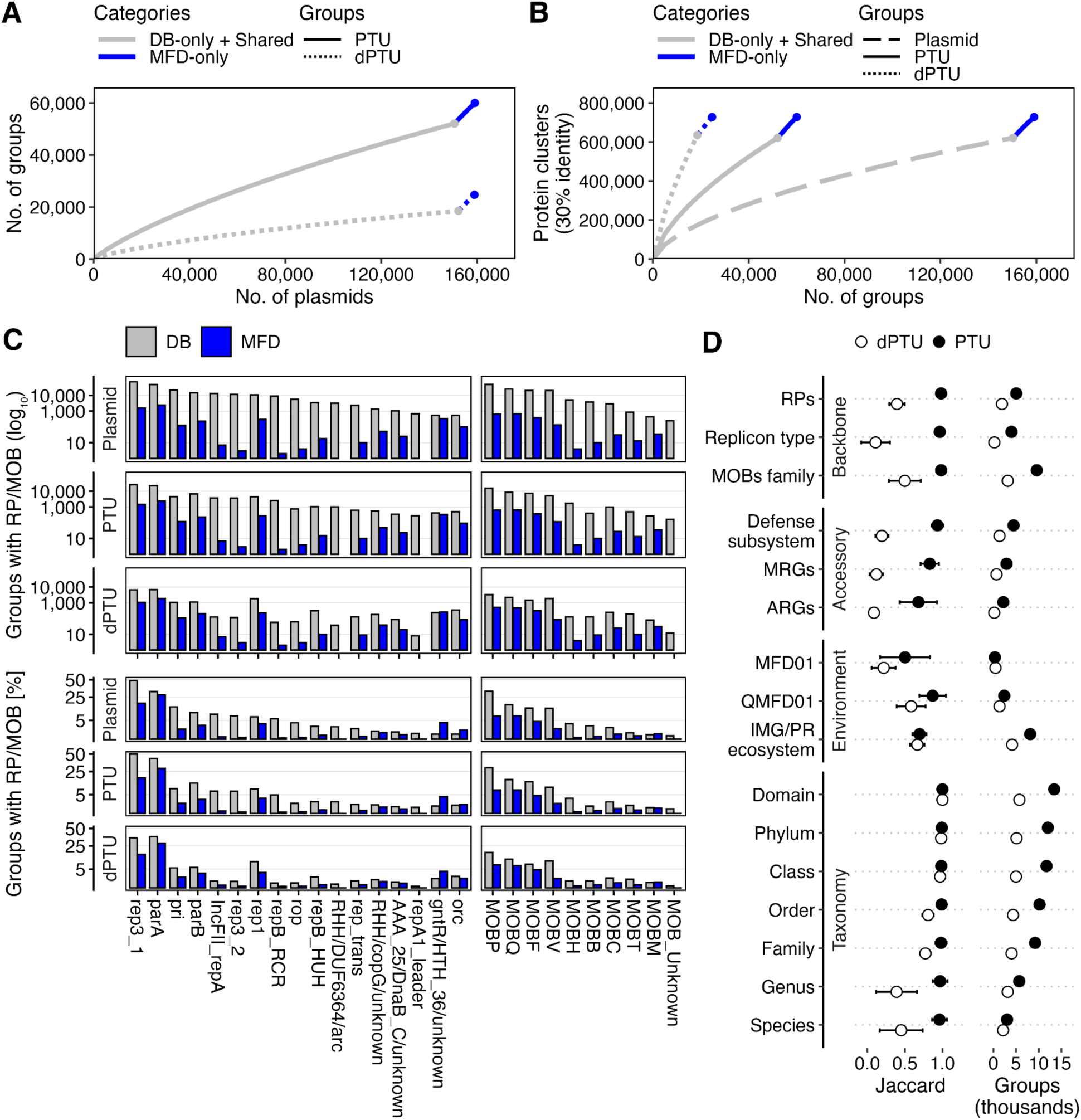
Danish terrestrial plasmids span diverse taxonomic units and replicon families. **(A)** Rarefaction curves showing the number of PTUs and distant PTUs (dPTUs) against the number of plasmids. **(B)** The number of plasmids, PTUs and dPTUs against the number of protein clusters at 30% amino acid identity. **(C)** Prevalence of plasmids with a given marker gene. The left facet represents the replication-associated proteins (RPs) while the right facet delineates the mobilases. **(D)** Biological coherence of PTUs vs. dPTUs across different traits measured using weighted Jaccard similarities excluding singletons **(Methods)**. Error bars indicate the weighted standard deviation across groups, with weights based on the number of plasmid pairs per group. Groups refer to the number of groups in the calculation in thousands.

Similarly, only 36 MFD plasmids could be successfully typed by PlasmidFinder^9^, consistent with its focus on clinically-relevant plasmids. To gain insight into replicon diversity and characterize the groups formed by our dPTU definition, we curated a set of RPs based on a diverse array of model replicons **(Methods; Supplementary Note 1;** **Fig. 2C**). Forward searching the final RP set **(Methods)**, we identified alignments to proteins encoded by 3,505 of 8,874 (40%) of MFD plasmids. Of these, 1,585 (45% of 3,505) were represented by Rep3_1 or Rep3_2. By contrast, these 17 RP families spanned 122,937 of 150,308 (82%) DB plasmids, of which 75,033 (61%) were represented by Rep3_1 or Rep3_2. Homologs of at least one of the 17 RPs could be identified on 40,740 of 52,062 (78%) DB PTUs and 3,325 of 8,158 (41%) MFD PTUs. We used the RPs and other annotations (representing putative “traits”) to assess the biological coherence of our PTUs and dPTUs. Coherence was calculated separately for each trait using only plasmids with available annotations and groups containing at least two such plasmids **(Methods)**. Values are reported as weighted mean Jaccard similarity ± weighted standard deviation **(Fig. 2D)**. For selected traits, these values are followed by the number of contributing groups and plasmids in those groups, reported as (mean ± standard deviation; groups; plasmids) **Supplementary Fig. 5**.

We observed high coherence of PTUs for RPs (0.98 ± 0.05; 11,789; 94,292), replicon type (0.96 ± 0.07; 4,009; 39,071) and MOB family (0.98 ± 0.04; 9,577; 84,269), indicating that the PTUs capture plasmids sharing a conserved backbone (replication and mobilization machinery) **(Fig. 2D; Supplementary Fig. 5)**. This coherence extended to inferred host taxonomy, with similarity remaining high from domain (1.00 ± 0.00) to species-level (0.96 ± 0.10; 3,036; 19,487). Taxonomy was assigned using MMseqs2 taxonomy^32^ with GTDB^33^ references, and coherence was calculated only for non-singleton groups with assignments at the respective taxonomic rank **(Methods)**. Because reference plasmids accounted for most high-resolution taxonomic assignments, especially at genus- and species-level, these values are dominated by the reference set; however, MFD plasmids that could be assigned at these ranks (819 of 8,744 at species-level) showed similarly high coherence (**Supplementary Fig. 6**). By contrast, PTU coherence was slightly lower for more variable accessory traits, including defense subsystems as defined in DefenseFinder^34–36^ (0.93 ± 0.08), metal resistance genes (MRGs) (0.83 ± 0.12) and antibiotic resistance genes (ARGs) (0.68 ± 0.25) **(Fig. 2D; Supplementary Fig. 5)**. For dPTUs, weaker conservation was observed with the RP (0.52 ± 0.21) and MOB family (0.50 ± 0.21) being the most coherent categories, indicating that plasmids grouped at this level may still retain some shared replication and mobility-associated traits. Host taxonomic coherence was also retained at broader ranks, most notably at the family-level (0.77 ± 0.07), but decreased at genus (0.39 ± 0.27) and species-level (0.45 ± 0.29) **(Fig. 2D; Supplementary Fig. 5)**. When compared to the Jaccard similarity for host-association at the PTU-level, the results suggest that the dPTUs as defined here reflect plasmid lineages that have sufficiently diversified to cover a broader taxonomic host range.

Notably, defense subsystems showed a level of PTU coherence (0.93 ± 0.08) that was closer to classical backbone traits, such as RPs (0.98 ± 0.05), replicon type (0.96 ± 0.07) and MOB family (0.98 ± 0.04), than to more variable accessory traits such as MRGs (0.83 ± 0.12) and ARGs (0.68 ± 0.25) **(Fig. 2D; Supplementary Fig. 5)**. By contrast, environment source was only weakly associated with PTUs and dPTUs. To assess whether plasmid groups were associated with similar environments, we quantified plasmid relative abundances across samples **(Methods)** and considered any quantification above zero as proof of presence. We refer to this quantification-based habitat preference as QMFD01, which displayed comparatively strong coherence in both PTUs and dPTUs (0.88 ± 0.18 and 0.60 ± 0.19, respectively). This indicates that at this environmental resolution closely related plasmids tend to share a common habitat origin. Taken together, these results support the idea that the PTUs capture conserved plasmid families with coherent backbone-associated traits (supported by coherence in taxonomy and lesser support of accessory genes), indicating biological coherence consistent with their proposed use as operational taxonomic units^15,17^. This is further supported indirectly by the coherence of inferred host taxonomy, while lower accessory coherence is consistent with variation in cargo expected within conserved plasmid families. The dPTUs group more distantly related plasmids through shared protein content and sequence similarity, serving as a more broad, yet coherent, biological unit for plasmid classification and future tracking.

The plasmid sequences displayed RP-specific GC content and length variation, consistent with differences in taxonomic origin and plasmid family origin **(Fig. 3)** and is discussed further in **Supplementary Note 1**. Interestingly, two clusters emerged from the curation process that were typified by PF01051 (Rep_3), a family characterized by a variable number of WH domains^18^. Despite sharing the same PFAM, no homology could be detected when performing remote sequence homology searches (likely due to the limits of local sequence alignments) and the plasmids represented by the two clusters differed substantially in terms of their length and co-occurring markers **(Fig 3A; Supplementary Fig. 7).** The two clusters (Rep3_1 and Rep3_2) were found on 29,989 and 11,297 dereplicated plasmids, respectively. Plasmids encoding these two clusters had median lengths of 18.8 kb and 142.3 kb, respectively (two-tailed Wilcoxon test, p<1e- 5), further suggesting that these are distinct protein families associated with distinct plasmid lineages. A total of 2,526 out of 11,297 Rep3_2 plasmids also encoded the IncFII_repA/repB2_RCR replicon associated proteins. IncF plasmids are known to be mosaic in nature, thus distinguishing individual RPs can shed light on the individual replicons^15^. Whereas naming conventions for IncF-type plasmid RPs reflect historical associations with one another, differentiating them based on sequence demonstrates again that IncF plasmids are formed from the combination of multiple, independently evolving plasmid lineages^37^. In sum, these results highlight that the RPs curated here may serve as useful hallmarks for future plasmid curation.

**Figure 3.**
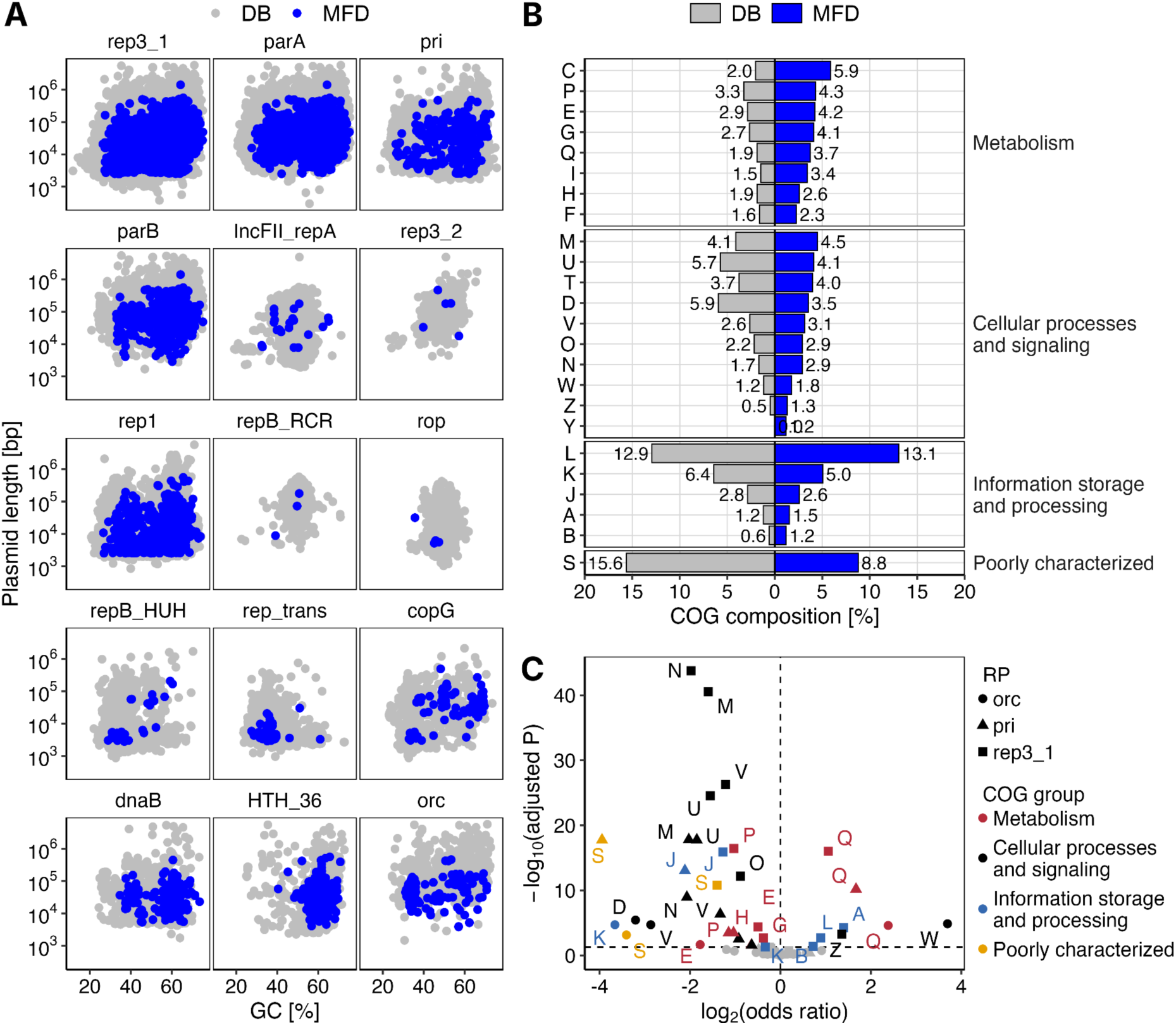
Danish plasmids span diverse replicon families and functions. **(A)** Sequence features of RP families colored by source (DB and MFD). Only RPs with hits to MFD plasmids are included, i.e., all except for repA1_leader and DUF6364. **(B)** Proportion of clusters of orthologous groups (COGs) assigned to the 30% identity protein clusters. **(C)** COG enrichment analysis comparing DB and MFD of three major RP families. The x-axis shows the log₂-transformed odds ratio (OR), with positive values indicating enrichment in MFD plasmids relative to DB plasmids. Thus, log2(OR) > 0 indicates enrichment in MFD relative to DB. The shapes refer to three selected RPs spanning diverse replicon types. The COG groups for metabolism are more highly represented in the MFD plasmids, notably Secondary metabolites biosynthesis, transport and catabolism (Q), which is enriched across all three RP families. Extracellular structures (W) and Cytoskeleton (Z) are also enriched in MFD plasmids carrying the origin recognition complex (orc) and rep3_1, respectively.

We next asked how MFD plasmids differed in functional content relative to their DB counterparts, focusing on three major RP families (Rep3_1, the most common RP; pri, a distinct primase-type RP associated with Bacillota plasmids; and orc associated with archaeal plasmids). These represent taxonomically and ecologically distinct plasmid families **(Supplementary Note 1).** While analysis of predicted protein clusters suggested enrichment in metabolism-related functions **(Figure 3B),** functional enrichment analysis is challenging because of differences in plasmid size, overrepresentation of mostly clonal plasmids represented by PTUs, and differences in annotation rates. To address this, we modeled the presence and absence of 20 COGs based on the consensus annotations of protein clusters from the plasmids **(Methods)**. We further included plasmid size and PTU as covariates in order to model enrichment (odds ratios) and assess confidence (**Supplementary Data 1).** This revealed that, across all three RPs, MFD plasmids were enriched for COG category Q (Secondary metabolites biosynthesis, transport, and catabolism), relative to their database equivalents. For Rep3_1, COG categories Q, Z (Cytoskeleton), and A (RNA processing) were the most enriched COGs. In addition, COG category W (Extracellular structures) was highly enriched in MFD orc plasmids. Examining the putative functions of COG category Q proteins in Rep3_1, pri, and orc-bearing plasmids suggested that frequent carriage of S-adenosine methyltransferases (SAMs) associated with diverse secondary metabolite-associated pathways was a driver across all three RP types. For Rep3_1 and pri, the most common PFAM of COG category Q proteins was adh_short_c2, a short-chain reductase (SDR) that is canonically associated with fatty acid biosynthesis. However, such SDRs are associated with a wide range of metabolic functions^38^. Interestingly, Rep3_1-associated COG category Z proteins included seven proteins with reference orthologs in Eukaryota, indicating potential HGT or a role in inter-kingdom interactions. This was further substantiated by the presence of eukaryotic-like protein domains, including HYR and ankyrin-repeats, among others **(Supplementary Data).**

### A novel *Methanosarcina*-associated plasmid lineage

Relatively little is known about the ecology of archaeal plasmids in the environment. Given this, and the observed enrichment of extracellular structure (COG category W) proteins in orc plasmids, we wanted to explore the potential role of MFD plasmids in host ecology across Denmark. Considering the relatively sparse representation of archaeal plasmids in the reference, we focused on the diversity, putative hosts, and ecological context of plasmids encoding the archaeal-type RP orc. Confirming the association with archaea, the orc RP was detected across 454 of 564 (83%) of the filtered plasmids with cultivated archaeal hosts. Amongst these, 126 were from MFD, and all of them were assigned MFD-only PTUs. This equates to a 20% increase in archaeal plasmids relative to the 627 from the reference collection. Almost all of the orc-bearing plasmids were assigned to the phylum *Halobacteriota*, including every MFD plasmid and all but five of the reference plasmids. Assessing the taxonomic assignments at greater resolution, 103 out of 120 plasmids in the new lineage were assigned to the genus *Methanosarcina*. Of the remaining 17, six formed a separate subclade of the Bog-38 genus, while 11 were assigned only to higher taxonomic ranks. Interestingly, the MFD and reference plasmids differed notably in GC content, averaging 36.9% and 58.2%, respectively. This is concordant with the difference in GC content between GTDB representatives of the classes *Halobacteria* and *Methanosarcinia* **(Supplementary Fig. 8)**. In the orc RP phylogeny, the MFD plasmids also comprised a distinct cluster with 120 of the 126 archaeal plasmids clustering together on the tree **(Fig. 4)**. This assignment was further supported by the presence of two reference plasmids from cultivated *Methanosarcina* (NZ_CP009527.1 and NZ_CP009519.1). In sum, these results suggest that the MFD plasmids include a novel lineage of *Methanosarcina*-associated plasmids **(Fig. 4)**.

**Figure 4:**
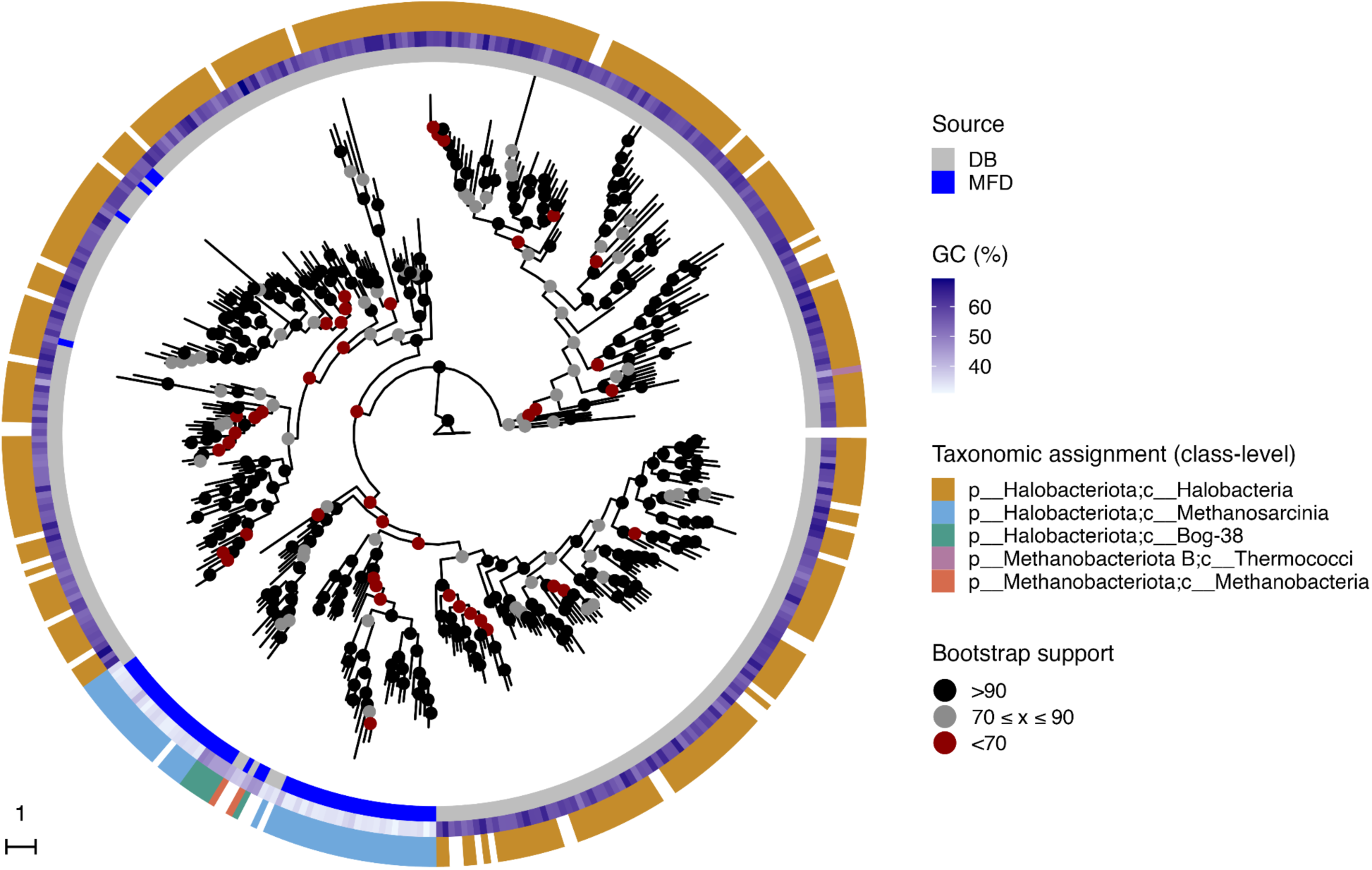
Mid-point rooted maximum-likelihood phylogenetic tree of the archaeal-type RP orc. Branch lengths were square-root transformed for visualization. Internal node points indicate bootstrap support. Heatmap grids denote, from left to right, the database source (DB or MFD), GC content (%), and putative taxonomic class of the plasmid host. The branch-length scale is shown in the bottom-left corner.

**Figure 5:**
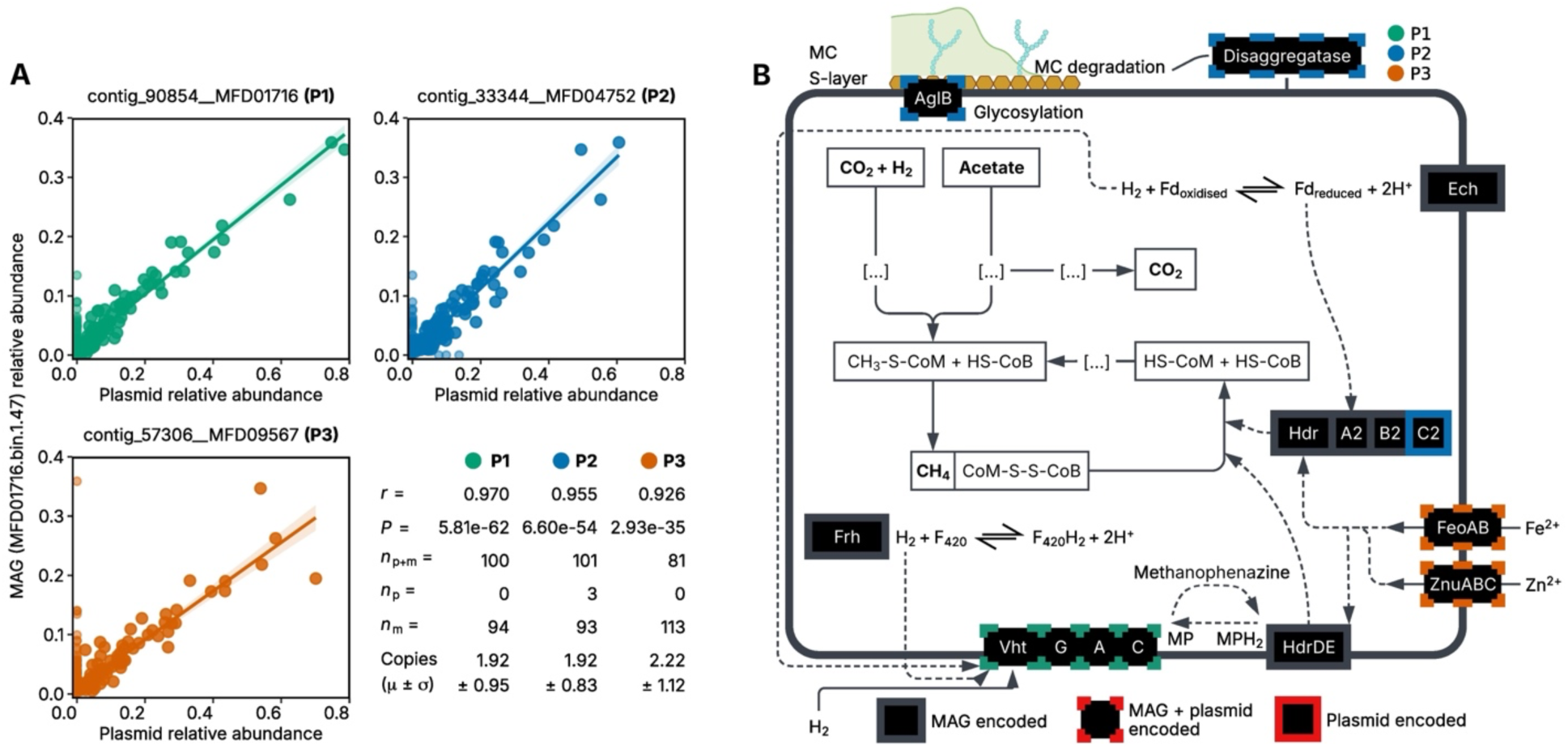
A) Abundance-based correlation of three *Methanosarcina* plasmids to the *Methanosarcina* MAG MFD01716.bin.1.47. Each point is one metagenome sample, plotting the per-sample relative abundance of a putative *Methanosarcina* plasmid (x-axis) against the relative abundance of the MAG MFD01716.bin.1.47 (y-axis). Each colored line is a linear regression with its 95% confidence interval (shaded), fitted only to samples where both the plasmid and the MAG were detected. In the inset table, *r* is the Pearson correlation coefficient between plasmid and MAG relative abundances across samples in which both were detected, and *P* represents the corresponding two-sided *P* value. The sample counts are *n*ₚhₘ for samples where both were detected, *n*ₚ for samples where only the plasmid was detected, and *n*ₘ for samples where only the MAG was detected. Copies (μ ± σ) is the mean ± standard deviation of the per-sample plasmid to MAG relative abundance ratio over the *n*ₚhₘ detected samples. **(B)** Simplified schematic of the predicted roles of proteins encoded by MAG MFD01716.bin.1.47 and three putatively associated plasmids (P1, P2, and P3) in methanogenesis and selected accessory functions. Established methanogenesis reactions follow Mand and Metcalf^40^, with Vht-associated hydrogen cycling and Heterodisulfide reduction (Hdr) following Kulkarni *et al*.^41^ and Yan *et al*.^42^, respectively. Only functions discussed in the text are drawn, and dotted lines indicate the earlier steps of each pathway, not shown. Selected redox carriers are shown, but energy conservation is omitted. Solid arrows trace the main pathway; dashed arrows indicate established or proposed functional connections involving the highlighted proteins. Unfilled boxes are metabolites, and bold ones cross the membrane. Black boxes with grey outlines indicate functions encoded by the MAG. Dashed colored outlines indicate functions also found on the MAG, while solid colored outlines indicate plasmid only functions (green for P1, blue for P2 and orange for P3).

To investigate the ecology of these plasmids, we identified potential hosts among the *Methanosarcina* metagenome assembled genomes (MAGs) from the MFD dataset^39^. Seven of the orc plasmids were highly correlated with a single high-quality *Methanosarcina* MAG, MFD01716.bin.1.47 **(Supplementary Table 2; Supplementary Fig. 9)**. We focused on three plasmids that complemented the metabolism (methanogenesis) and lifestyle of *Methanosarcina* **(Supplementary Note 2)** referred to herein as P1 (contig_90854 MFD01716), P2 (contig_33344 MFD04752) and P3 (contig_57306 MFD09567). All three were assembled from the same sample as the putative *Methanosarcina* host, but near-identical clonal plasmids (dereplicated at 95% sequence identity and 99% bidirectional coverage) were also recovered from multiple samples (two for P1, four for P2 and P3). The contig identifiers refer to representative plasmids. The relative abundance of the three plasmids showed strong correlations with the abundance of the MAG (two-sided Pearson tests against zero correlation, using samples where both were detected), with correlation coefficients of 0.970 for P1 (P = 5.81 x 10e-62; n = 100), 0.955 for P2 (P = 6.60 x 10e-54; n = 101) and 0.926 for P3 (P = 2.93 x 10e-35; n = 81). The estimated copy number was approximately two copies per genome for all three plasmids **(Fig. 5A)**.

Analysis of the putative functions encoded on the three plasmids suggested convergent and complementary adaptations to the plasmids’ source habitats, which were chiefly sphagnum acidic bogs **(Fig. 5B; Supplementary Fig. 10)**. Sphagnum acidic bogs are anaerobic freshwater wetlands that accumulate acetate from Sphagnum degradation, providing ideal conditions for acetate-utilising Methanosarcinaceae^43,44^. As expected, MFD01716.bin.1.47 encodes all steps necessary for acetoclastic methanogenesis **(Supplementary Table 6; Supplementary Table 7)**. Interestingly, two of the three plasmids also encode functions linked to the regeneration of cofactors associated with the terminal step of methanogenesis^40^ **(Fig. 5B)**. Namely, P1 encodes a copy of VhtGAC hydrogenase (which supports regeneration of coenzymes B and M upstream of membrane encoded HdrDE through hydrogen cycling), with 60% amino-acid identity to the chromosomally-encoded copy^41^ (**Fig. 5B; Supplementary Table 3**). P2 encodes heterodisulfide reductase subunit C HdrC, completing the reductase complex (HdrA2B2C2) whose remaining subunits, HdrA2 and HdrB2, are encoded on the chromosome **(Fig. 5B; Supplementary Table 4)**. This complex likewise regenerates coenzymes B and M^42^. In addition to complementing cofactor regeneration by encoding HdrC, P2 also encodes plasmidic copies of a putative AglB/STT3-family oligosaccharyltransferase and a disaggregatase-related protein, the latter sharing 60% amino-acid identity with their chromosomal counterpart **(Fig. 5B)**. These proteins are associated with cell surface features, including methanochondroitin, a specialized extracellular matrix that facilitates aggregation of *Methanosarcina*^45–47^. AglB is required for surface-attached growth in *Methanococcus maripaludis*^45^, and disaggregatase promotes dispersal of *M. mazei* aggregates^48^. The plasmid-encoded homologs may therefore influence transitions between aggregated and dispersed growth^46^, contributing to the host’s adaptation to changing conditions **(Fig. 5B)**. The presence of alternative copies of genes linked to this process is intriguing, and may result in heterogeneous cell surface and attachment properties in *Methanosarcina* populations. These genes could also facilitate horizontal plasmid transfer by altering cell attachment and dispersal, and may together contribute to host ecological adaptability by facilitating transitions between aggregated and dispersed growth. Finally, P3 encodes an iron transporter operon *feoAB* and the zinc transporter operon *znuABC* **(Fig. 5B; Supplementary Table 5)**. The abundance of these micronutrients is often a key limit on microbial growth in oligotrophic environments, and the role of plasmids in conferring nutrient scavenging properties is well established^49–52^.

Our observations in *Methanosarcina* are in line with other recent studies reporting on plasmid ecology. Plasmid-encoded copies of chromosomal genes have been increasingly observed in environmental plasmids^53^, as we observed for VhtGAC and the disaggregatase-related protein. These alternative copies could provide yet unknown fitness benefits such as increased gene dosage or functional diversification. Such flexibility may support adaptation to changing environmental conditions while preserving the chromosome-encoded function. For example, increased gene dosage of VhtGAC could enhance its native role in hydrogen cycling, promoting cofactor regeneration under competitive conditions^41^. Alternatively, it could represent a specialized variant that shows better enzymatic activity under differing environmental/metabolic conditions. Furthermore, we identified the remaining HdrA2B2C2 subunit as putatively plasmid encoded, highlighting the potential role of chromosome-plasmid complementarity in prokaryotic evolution. This complementarity could serve as an adaptive mechanism to facilitate the native reductase complex activity in the absence of a functional copy of HdrC. Simultaneously, it could serve as an additional mechanism facilitating plasmid maintenance across generations. The presence of this gene on a plasmid may also suggest it is linked to a trait that is dispensable depending on the prevailing environmental conditions. Regardless of the exact mechanism, these results illustrate a potentially important limitation of inferring cellular metabolism from host chromosomes alone, and the necessity of considering the mobilome to determine the host’s metabolic and ecological potential.

## Discussion

The publicly available collection of known plasmids is skewed towards clinical isolates and model organisms. It follows that domain-specific knowledge of plasmids, such as their nomenclature, grouping and functional characterization, has been historically biased towards a limited representation of prokaryotic diversity. Here, largely through deep long-read sequencing, we introduce a new set of 8,744 curated, putatively complete environmental plasmids that increase known plasmid diversity at the PTU-level by 15% relative to existing resources (**Figure 1**). Considering the historical characterization of plasmids covered only a fraction of the ones deposited in public databases, we took the opportunity to reassess hallmark plasmid genes in light of increased environmental representation. We curated 17 RP families to function as plasmid markers, covering 82% of plasmids in the public domain and 40% of our set. In the process, we incorporated the recent work of Nishimura *et al*.^18^, which focused on the winged-helix domain, which could characterize 57% plasmids from the PLSDB dataset, and generalized it to multiple known and proposed RP families. Importantly, although up to 82% of plasmids had an identifiable RP homolog, homology alone is insufficient evidence to assert that two proteins retain the same function. Thus, we view this RP set as a first step towards establishing a comprehensive RP plasmid hallmark repertoire that, in tandem with robust topological inference, can confidently identify complete plasmid sequences in the environment.

Beyond near-identical clones, PTUs group plasmids by overall DNA similarity, resolving sequence relationships that replication markers alone cannot^13,15,17^. PTUs follow the rationale that similar DNA sequences have diverged recently and mirror the rule of thumb for ANI-based species definition in bacteria, despite using different ANI and AF thresholds. Furthermore, PTUs have been shown to have a coherent replicon typing and host taxonomy (either as single-host or promiscuous), corroborating them as useful biological units^15^. Here, taking inspiration from the AAI sequence similarity to infer taxonomic group at the genus-level for bacteria^30,31^, we proposed the use of the same technique to identify distant PTUs (dPTUs). We demonstrated that the findings of PTU coherence are generalizable to strict habitat preference and ancillary gene content (e.g., MRG, ARG), while dPTUs group more distantly with less narrow yet coherent traits (**Figure 2**). Plasmids contribute to genome evolution through both gene acquisition and diversification. They bring a flexible set of genes to the genome, potentially complementing or expanding the chromosomal set. Historically, much of our knowledge of plasmid gene content has come from cultivated isolates, particularly from clinical settings, where antimicrobial resistance has been a major focus. Environmental plasmids remain less represented in reference collections, and their functional potential underexplored. We used COG categories to compare the broad functionality of terrestrial environmental and reference plasmids carrying a curated set of three relevant RPs. We observed that the Danish plasmid dataset is enriched in a focused subset of COG categories pointing to their environmental roles. For example, *Methanosarcina* carry plasmids that augment their central metabolism (methanogenesis) and help them to scavenge micronutrients. This is particularly notable in a genus whose acetoclastic metabolism was itself shaped by horizontal gene acquisition^20,21^. These findings illustrate the high potential for the discovery of novel biology in archaea, whose plasmids remain poorly represented despite their hosts’ importance in biogeochemical cycling.

Our set of 8,744 terrestrial plasmids obtained from the relatively small geographical area of Denmark, resulted in a substantial increase in known plasmid diversity and provides fresh perspective on historical challenges such as plasmid grouping and functional contribution to the genome starting from a wider and more diverse base. Our results suggest that plasmid biology remains greatly underexplored and that outstanding degrees of diversity remain to be discovered in terrestrial microbiomes.

## Methods

### Plasmid prediction

We collected the assembled contigs from the Microflora Danica (MFD) project^39,54^, a national-scale metagenomic survey of Danish environmental microbiomes spanning soil, sediment, and water habitats across Denmark. Plasmid prediction was performed on ∼330.9 Gbp of short-read (Illumina) and 295.7 Gbp of long-read (ONT) contig assembly data, representing assemblies from 9,972 Illumina-sequenced metagenomes^54^ and 154 ONT-sequenced metagenomes^39^ collected across all regions of Denmark (**Fig. 1)**. The Illumina assemblies were obtained from a pre-publication version of the MFD dataset and therefore comprise fewer metagenomes than the published release. These metagenomes were derived from soil (Illumina, n = 7,970; ONT, n = 25), sediment (Illumina, n = 1,464; ONT, n = 28), and water (Illumina, n = 267; ONT, n = 1) samples. The plasmid prediction was carried out using four state-of-the-art plasmid prediction tools: geNomad^25^ (v1.7.4), PLASMe^27^ (v1.1), Plasmer^26^ (v23.04.20), and viralVerify^28^ (v1.1); all run with default settings. In total, 6,388,240 Illumina and 1,561,754 ONT contigs were predicted to be plasmidial.

### Topology classification by read-mapping

Plasmidial contigs were classified as circular, linear or unresolved by mapping sequencing reads back to the assembly from which each contig was derived. Reads were mapped with minimap2^55^ (v2.28-r1283) using -cx lr:hq --secondary=no for ONT data and -cx sr --secondary=no for Illumina data, retaining only primary alignments. Supporting alignments were retained at ≥95% identity across ≥99% of the read length for ONT reads longer than 1,000 bp, and at ≥99% identity for Illumina contigs longer than 2,500 bp with ≥5x coverage; reads and contigs below these thresholds were discarded.

Circularity was predicted from reads spanning the artificial breakpoint introduced during assembly. For ONT contigs this required ≥1,000 bp of continuous alignment across the two contig ends, with ≥500 bp of support on either side of the breakpoint. For Illumina contigs it required at least three read pairs bridging the two ends in reverse-forward orientation, mapping within 201 bp of a terminus (151 bp read length plus 50 bp). Linearity required read support at both contig ends, with no terminal overhang exceeding 5 bp for ONT contigs or 1 bp for Illumina contigs. The assessment of circular topology from long-read data was implemented in circure (https://github.com/bertils/circure), which we made for this purpose.

### Databases and their curation

To place the MFD plasmids in the context of the known plasmids, we compiled a reference collection of plasmids from PLSDB^11^ (v2024_05_31_v2; n = 72,556) and IMG/PR^12^ (downloaded in 2024; n = 154,680). Because we only included putatively complete plasmids from the MFD dataset in our analyses, we restricted the IMG/PR set to entries annotated as putatively complete.

The DB plasmids and the MFD plasmids were dereplicated with MMseqs2^56^ using 99% bidirectional coverage and 95% sequence identity (v17.b804f; --cluster-mode 2 --cov-mode 0 -c 0.99 --max-seq-len 20000000 --min-seq-id 0.95 --seq-id-mode 0 --kmer-per-seq-scale 0.5 -- alignment-mode 3 --wrapped-scoring 1). After dereplication, the MFD set was reduced from 13,287 to 12,504 plasmids, and the DB set from 227,246 to 164,816 plasmids **(Fig. 1)**. Throughout this paper, we only include and compare sequences that are putatively complete, which we refer to as plasmids; otherwise, we refer to (incomplete) sequences predicted to be plasmids as plasmidial contigs.

### Scoring plasmid sequences

The final scoring system included consideration of topology, source database, and functional content and was partially adapted from the system used to score IMG/PR plasmids^12^. Briefly, presence of a plasmid hallmark in mobileOG-db v.2.0^57^ (detected via diamond blastp --id 30 -e 0.001) gave a +3 to the overall score; the presence of a mobilase or mating pair formation (MPF) protein (detected via diamond blastp against the mobsuite database --min-seq-id 40 -e 0.001) added a +2 to the overall score; a circular topology supported by either the presence of direct terminal repeats for database-derived plasmids or read-mapping for ONT-derived MFD plasmids (see section “Topology classification by read-mapping”) gave a +1 for a maximum score of 6. Proteins annotated as mobilases were excluded from the plasmid hallmark screen when using mobileOG-db in order to avoid double counting.

A major component of the curation was protein database mobileOG-db^57^. For example, mobileOG-db differentiates serine resolvases to a separate category of recombination/integration/excision whereas geNomad, which identified hallmarks based on class-wise protein cluster enrichment, frequently identified clusters of serine resolvases associated with plasmids. Serine resolvases are associated with a diversity of MGEs and are not plasmid specific, thus encoding a serine resolvase does not qualify a contig as a plasmid alone^58^. Although part of this discarded fraction may represent non-plasmid sequences, some genuine plasmids may also have been removed if they did not encode a hallmark gene similar to ones derived from canonical plasmid families **(Fig. 1)**. Throughout this paper, when we write “DB” (or “Database”), we mean this specific reference collection that has gone through dereplication and curation **(Fig. 1)**.

### PTU inference

In order place the MFD plasmidome in the context of the known plasmid diversity, we curated and dereplicated set of 8,744 MFD plasmids together with 150,308 complete reference plasmids from IMG/PR and PLSDB, and grouped all plasmids into plasmid taxonomic units (PTUs) based on all-versus-all sequence similarity. All-versus-all alignment was performed using skani^59^ (v0.2.2; triangle --diagonal --sparse -m 50 -c 5) with sensitive parameters (-m 50 -c 5) to improve alignment accuracy for the relatively short sequences typical of a small plasmid **(Supplementary Fig. 11)**.

The resulting pairwise alignments were used to construct a sequence similarity network, in which edge weights between plasmid pairs were computed as the equally weighted product of ANI and the aligned fraction of the shorter plasmid, following the approach of Camargo *et al*. 2023^12^, retaining only pairs meeting a minimum threshold of 70% ANI and 50% AF in accordance with the PTU definition of Redondo-Salvo *et al*.^17^. PTUs were inferred using Leiden community detection^60^ as implemented in pyLeiden (v0.1.12; -n 5, -r 0.9; https://github.com/apcamargo/pyleiden). The Leiden resolution parameter (-r parameter) controls how finely the network is split (lower values give fewer, larger groups, whereas higher values give more, smaller groups). We chose a resolution of 0.9 as it represented the point of diminishing returns, where further increases in resolution broke existing multi-member PTUs into singletons rather than forming new ones **(Supplementary Fig. 12)**.

To compare the MFD plasmidome with the Global Soil Plasmidome Resource (GSPR)^4^, we took the putatively complete plasmids from GSPR v3.0 that were not already in our reference collection and added them to the dereplicated and curated plasmid set (MFD + reference collection). We then inferred PTUs on this combined dataset using the same workflow as above (skani all-versus-all alignment, a similarity network thresholded at 70% ANI and 50% AF of the shortest plasmid, and Leiden clustering with -n 5 -r 0.9; see "PTU inference"). For the comparison we kept only PTUs that contained at least one GSPR plasmid.

### dPTU inference

Plasmids were clustered into distant plasmid taxonomic units (dPTUs) by calculating the average amino acid identity (AAI) and the percentage of conserved proteins (POCP) between the plasmids using CompareM^61^ (v0.1.2) with the following settings: “--evalue 0.00001”, “--per_identity 40”, “-- per_aln_len 50”, “--sensitive”. Cases of higher than 50% POCP and over 70% AAI were selected and used to construct an undirected graph with edges weighted by AAI, using the igraph package^62^ (v2.1.4) in R^63^ v4.4.1. The graph was then clustered using the Leiden algorithm^60^ with a resolution of 0.95, 5 iterations, and incorporating the edge weights.

### Annotation of plasmid proteins

Open reading frames were predicted for the plasmid sets using Prodigal^64^ v2.6.3, and eggNOG- mapper^65^ v2.1.12 with default settings was used for functional annotation of proteins (eggNOG Database^66^ v5.0.2; DIAMOND^67^ v2.0.11; MMseqs2^56^ v13.45111). CARD^68^ v3.4 was used as the reference database for ARG annotations using diamond blastp v2.0.11 with a minimum identity filter of 90% and e-value < 1e-5. Results were filtered using a manually curated index file of CARD to remove resistance derived from mutations to housekeeping genes, to delineate between efflux/membrane permeability resistance mechanisms and to provide major drug class assignments. Plasmid sequences were screened for defense and anti-defense systems using DefenseFinder^34–36^ (v2.01; -a). Plasmid replicon typing was annotated with PlasmidFinder^9^ (v2.1.6; -l 0.20 -t 0.50 -x).

Protein clusters were predicted using MMseqs2^56^ v.2.0.11 easy-cluster (mmseqs easy-cluster -i 0.3 -c 0.5). Metal resistance genes (MRGs) were annotated using the predicted and experimental sequences from BacMet^69^ v2.0 with diamond blastp (minimum sequence identity 40% and e-value < 1e-5). These results were filtered to remove transcriptional regulators by removing hits with related keywords in the header.

### RP functional enrichment analysis

To reduce bias from uneven annotation depth between DB and MFD plasmids, COG categories were assigned at the level of ∼30% amino acid identity clusters (described above) rather than individual proteins. For each cluster, the proportion of annotated members assigned to each COG category was calculated, counting multi-category assignments toward every category listed. A category was accepted as part of a cluster’s consensus annotation if it was supported by at least 25% of annotated members, in clusters with at least three annotated members. Every protein in a cluster inherited the cluster’s consensus categories, and a plasmid was scored as carrying a COG category if at least one of its proteins belonged to a cluster with that category in its consensus annotation.

For the three RPs examined, the presence or absence of each COG category was modeled by logistic regression with source (MFD vs. reference collection) as the predictor and log-transformed protein count as a covariate to account for plasmid size. Because closely related plasmids are not independent observations, standard errors were clustered by PTU. Models were fitted with the fixest package^70^ v0.14.2 in R^63^ v.4.5.3. Odds ratios are reported with Wald 95% confidence intervals, and P values were corrected for multiple testing across COG categories using the Benjamini-Hochberg procedure^71^. Categories with fewer than five plasmids in any source x presence combination were excluded from testing. Odds ratios above 1 indicate enrichment in MFD plasmids relative to reference database plasmids at equivalent plasmid size.

### Rarefaction analyses of plasmid and protein-cluster diversity

Rarefaction was calculated in R^63^ (v.4.3.2) using vegan^72^ (v2.7.1). Expected PTU and dPTU richness was calculated from plasmid counts using *rarecurve*. Expected protein-cluster richness was calculated from the number of plasmids, PTUs or dPTUs containing each protein cluster by summing detection probabilities using *drarefy*. Sampling was without replacement, using steps of 100 for PTU and dPTU rarefaction and 5,000, PTUs or dPTUs for protein-cluster rarefaction. DB-only + Shared categories were included first, followed by MFD-only. Protein clusters already found in the DB-only + Shared category were not counted again when adding the MFD-only.

### sylph quantification

The relative abundance of the species-level MAGs and plasmids was computed using sylph^73^ v0.6.1. Briefly, the short-read metagenomic samples from the Microflora Danica collection^54^ were sketched using the sketch command. The species-level representative genomes from Denmark were retrieved from two different studies^39,54^ and dereplicated using dRep^74^ (v.2.6.2; -comp 50 - con 10 -sa 0.95 -nc 0.4). Within each cluster, representative priority was given, in order, to cMAGs, MIMAG high-quality MAGs and long-read MAGs, and the resulting genome representatives were augmented with the plasmid set identified in this study. The reference sequences (genomes and plasmids) were sketched using the sketch command and the sample sketches were profiles using the profile command with min_kmers 20 and –read-seq-id 99.5 option.

### Taxonomic assignment of plasmids using MMseqs2

Putative host associations were assigned for plasmid sequences using MMseqs2 taxonomy^32^ (mmseqs taxonomy --tax-lineage 1 --majority 0.5 --vote-mode 1 --lca-mode 3 --orf-filter 1) and mmseqs createtsv. The reference database used was GTDB^33^ v220.

### Plasmid replication protein curation

RPs were identified using a combination of reference databases, manual curation, and iterative sequence alignment **(Supplementary Note 1)**. Briefly, an initial database of RP protein sequences was compiled by combining the database from Nishimura *et al*.^18^ with all entries in mobileOG-db^57^ that either aligned with a protein from Nishimura *et al*.^18^, or which were annotated as an RP based on the gene name and/or major and minor mobileOG categories. The merged protein set was aligned against Swiss-Prot^75^ (release 2025_01) using diamond (--id 40 --query-cover 80), and the aligned Swiss-Prot entries were manually curated to reduce the initial database to a subset of RP families. All plasmid proteins were then searched against this set (diamond blastp --ultra-sensitive) to curate a final set of potential RPs. RPs were then clustered at the level of protein family as follows: the sequences were clustered using mmseqs --min-seq-id 30 --cov- id 50 --cov-mode 0 -s 7, then the representative sequences were aligned to one another using diamond blastp --ultra-sensitive, and any degree of sequence homology was taken as evidence to collapse the cluster. The Levenstein distance of each cluster member was calculated relative to the cluster representative and if the cluster member differed by more than 4 standard deviations above the cluster mean, the cluster member was removed before multiple sequence alignment. The final set was then subject to an alignment parameter sweep to evaluate within-family variation **(Supplementary Note 1)** and set the final search cut-off of 40% identity and 50% query coverage, equivalent to the Enzyme Commission’s recommendation for transferring orthology assignments between folds^76^. For the orc tree, representative sequences for the orc family were extracted from the final RP set **(Supplementary Note 1)** and aligned using FAMSA^77^ with default settings and final tree prediction using IQ-TREE3^78^ (iqtree3 --alrt 1000 -B 1000 --boot-trees), including model selection.

### Biological coherence of PTUs and dPTUs

Trait similarity was calculated separately for PTUs and dPTUs in R^63^ (version 4.3.2). For each trait, annotations were converted to sets of unique labels. Pairwise Jaccard similarity was calculated within each group (PTU or dPTU) using a custom R function, jaccard(), as the number of shared labels divided by the number of labels in either plasmid (intersect() and union() in base R). The mean pairwise similarity was used as the Jaccard similarity per group for a trait, reflecting the extent to which plasmids in the same PTU or dPTU shared that indicated trait. The overall scores were then calculated as the weighted mean of these scores, with weights corresponding to the number of plasmid pairs in each group. The weighted standard deviation of was calculated using the number of plasmid pairs in each group as weights. Missing values were excluded from the calculations and could therefore not inflate the Jaccard score as shared absences. RPs refers to the curated replication-associated protein families described in the section “Plasmid replication protein phylogenies”; Replicon type to PlasmidFinder assignments (see section “Annotation of plasmid proteins”); MOB family to relaxase assignments obtained using the MOB-suite database (see section “Scoring plasmid sequences”); defence subsystem to DefenseFinder output (see section “Annotation of plasmid proteins”); MRGs to BacMet annotations (see section “Annotation of plasmid proteins”); and ARGs to CARD annotations (see section “Annotation of plasmid proteins”). The habitat descriptor MFD01 was derived from the MFD sample ontology for the source assembly, whereas QMFD01 was inferred from sylph-based profiling across samples (see section “sylph quantification”). IMG/PR ecosystem labels were obtained from the IMG/PR metadata. Taxonomic labels were taken from the MMseqs2 assignments described in “Taxonomic assignment of plasmids using MMseqs2”; root assignments were excluded **(Fig. 2D)**.

#### Stop-codon recoding in the putative *Methanosarcina* lineage

Contigs assigned to the putative *Methanosarcina* lineage (**Fig. 4**) were re-annotated using an alternative genetic code in which the amber codon (UAG) is read as pyrrolysine rather than as a translational stop^79^ (translation table 34). Under the standard bacterial and archaeal code (table 11), open reading frames containing in-frame UAG codons, a well documented feature of Methanosarcinaceae. Open reading frames for these contigs were therefore predicted with Prodigal^64^ specifying the translation table as -g 34 (v2.6.3), which terminates genes only at UAA and UGA and reports in-frame UAG positions as an ambiguous residue (X) in the translated sequence. The resulting protein sequences replaced the standard-code predictions for these contigs in all downstream annotation steps (see "Annotation of plasmid proteins").

### Comparison of plasmid and host genome gene content

The open reading frames (ORFs) were predicted for MAG MFD01716.bin.1.47 with Prodigal v2.6.3 specifying -g 34 and otherwise default parameters, as described above, and submitted to BlastKOALA^80^ for KEGG Orthology (KO) assignment **(Supplementary Table 6)**. The resulting KOs were compared with those assigned to the plasmid proteins by eggNOG-mapper (see "Annotation of plasmid proteins"). To capture chromosome-plasmid paralogs, the plasmid proteins were additionally searched against the MAG ORFs with MMseqs2 (v17.b804f; easy-rbh --cov-mode 1 -c 0.5 --min-seq-id 0.3), retaining reciprocal best hits. The putative AglB/STT3- family oligosaccharyltransferase encoded by P2 was annotated as STT3 by eggNOG-mapper. After reviewing its Pfam domains, we annotated it as an AglB/STT3-family oligosaccharyltransferase (AglB). Separately, BlastKOALA assigned an AglB-associated KO to a MAG-encoded protein, but no reciprocal best hit was recovered between the two proteins under the specified MMseqs2 criteria, suggesting that the plasmid and chromosome encoded distinct copies of AglB-like proteins.

### Co-abundance correlation of plasmids and MAGs

To identify putative plasmid-host associations, we co-correlated the relative abundance of MFD plasmids and metagenome-assembled genomes (MAGs) across samples using sylph^59^ as described in a previous section (see “sylph quantification”). We filtered for the putative Methanosarcina-associated plasmid lineage. For each pair of plasmid and MAG, we computed the Pearson correlation of their relative abundance across samples where both were present, with P values derived from a two-tailed t-test and corrected for multiple testing using the Benjamini-Hochberg false discovery rate correction^71^ **(Supplementary Table 2)**, all implemented in R^63^ (v4.3.2). We retained pairs detected together in at least 10 samples with a correlation coefficient greater than 0.5. These associations are listed in **Supplementary Table 2**, ordered by adjusted P value.

## Data availability

The metagenomic sequencing data and assemblies from the Microflora Danica project used in this study are available under accession PRJNA1071982 and PRJEB58634. The full set of plasmids analysed in this study are available on Zenodo at https://zenodo.org/uploads/22944851, together with the associated metadata, predicted ORFs, eggNOG-mapper annotations, GFF files and mobileOG-db hallmark-gene hits. The curated set of replication-associated proteins is available on Zenodo at https://zenodo.org/records/21099754.

## Code availability

The code used for the analyses and to generate the figures is available on GitHub at https://github.com/bertils/pfd. circure is available on GitHub at https://github.com/bertils/circure.

## Acknowledgements

The authors thank Eduardo Rocha for his feedback on the manuscript, and Jeppe Støtt Bøjer for discussions on methylation and putative host associations.

## Author information

Authors B. Sørensen and C. L. Brown contributed equally.

## Author contribution

B.S. and F.D. conceived the study. B.S., C.L.B., M.S. and F.D. designed the study. B.S. predicted and curated the plasmids with the input of C.L.B., C.L.B. predicted and curated the replication proteins of the plasmid and reconstructed their phylogeny, with input from P.H.. B.S. created the tool circure to predict completeness and topology of the putative plasmids and C.L.B scored them according to their reliability. B.S. and C.L.B. annotated the plasmids according to habitat origin, genetic potential and putative host. B.S. computed the PTUs and M.S. computed the dPTUs. B.S. analysed the coherence of plasmids annotation (traits) across PTUs and dPTUs. B.S. curated the assignment of the novel plasmid lineage to the genus *Methanosarcina* and reconstructed the putative integrated biology of the host together with its plasmids. B.S., C.L.B. and F.D. wrote the manuscript with input from all the authors.

## Ethics declaration

The authors declare no competing interests.

