## Supplementary Information for "Nation-scale recovery of complete environmental plasmids pushes the boundaries of plasmid diversity through long-read sequencing"

Bertil Sørensen[1^], Connor L. Brown[1^], Mantas Sereika[1], Søren Heidenbach[1], Philip Hugenholtz[1,2], Mads Albertsen[1], Francesco Delogu[1,3]*
[1]Center for Microbial Communities, Aalborg University, Denmark
[2]Australian Centre for Ecogenomics, School of Chemistry and Molecular Biosciences, The University of Queensland, Brisbane, Queensland, Australia
[3]Department of Computer Science, Aalborg University, Aalborg, Denmark
^Shared first authorship
*Corresponding author

### Supplementary Information

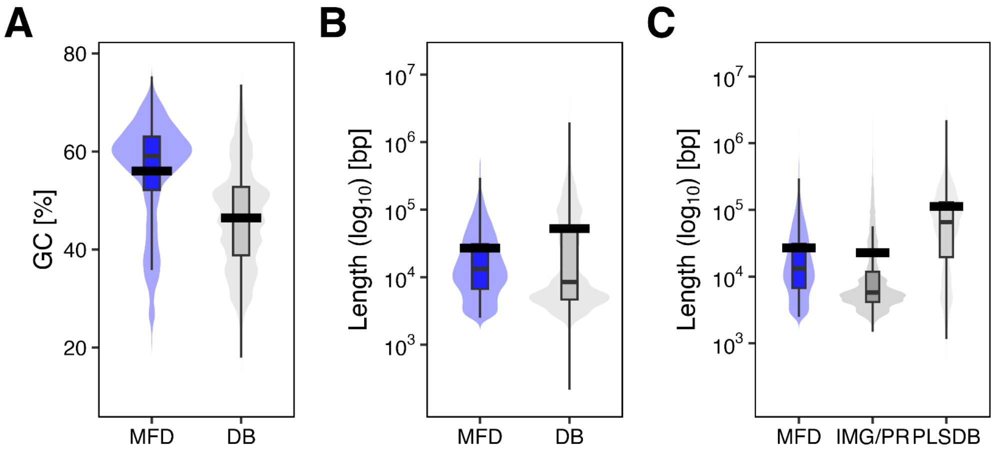


**Supplementary Figure 1.** Distribution of GC content in the plasmids from MFD and the reference collection (DB) **(A)**, length distribution for MFD and DB **(B)**, and length distribution for MFD, IMG/PR, and PLSDB **(C)**. Thick horizontal black bars indicate the mean. Box edges (hinges) show the 25th and 75th percentiles, and the line inside each box shows the median. Whiskers extend to the most extreme values within 1.5 times the interquartile range of the box edges. Outlier points are not shown.

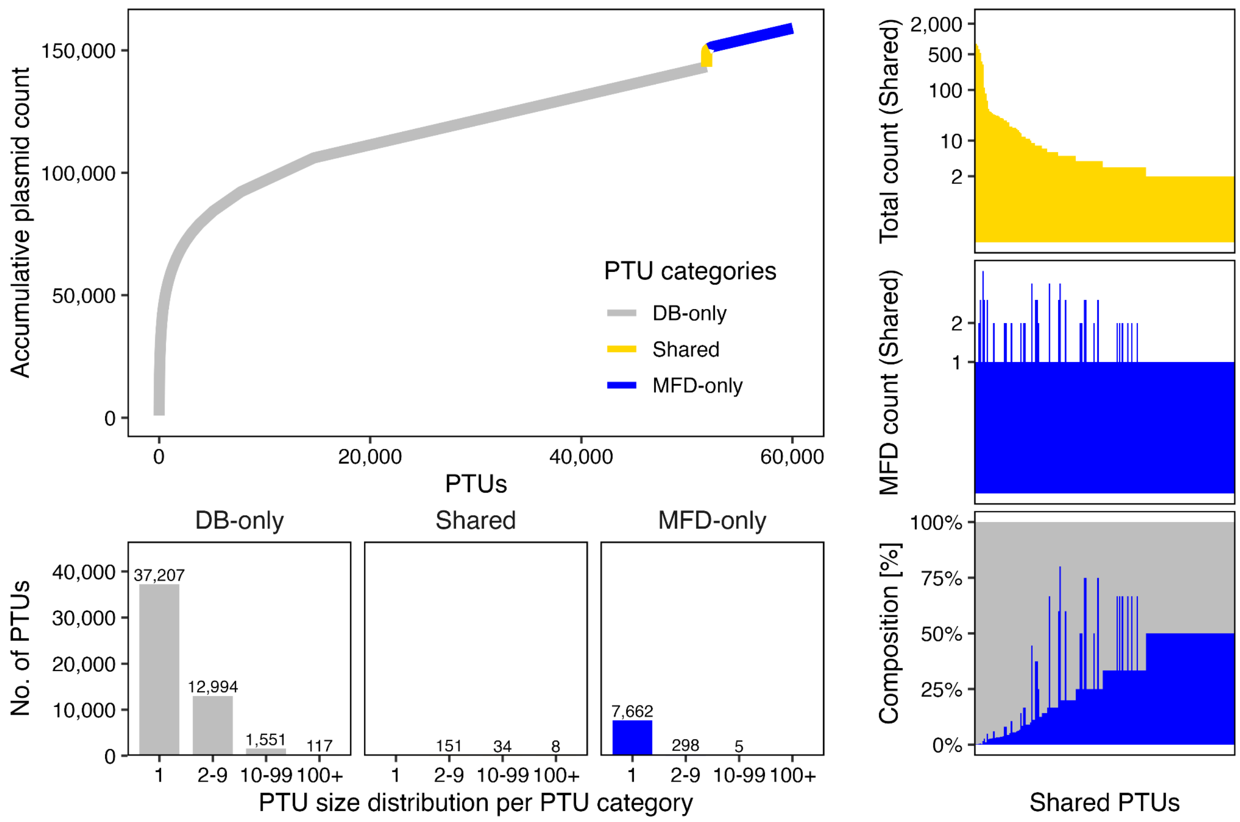


**Supplementary Figure 2.** Overview of the PTU inference of MFD and reference collection plasmids. The 8,744 MFD and 150,308 DB plasmids were clustered into 60,026 PTUs, each classified as DB-only, MFD-only, or Shared if a PTU contained plasmids from both sources. The accumulation curve (top left) plots the cumulative number of plasmids as PTUs are added in order of decreasing size. The bar plots (bottom left) give the number of MFD-only, DB-only and Shared PTUs within each size category. The three panels on the right give an overview of Shared PTUs, ordered from largest to smallest, showing the total count of plasmids per Shared PTU (top), the count of MFD plasmids in such Shared PTUs (middle) and the composition in percent of the number of plasmids from either source (bottom). Shared PTUs are dominated by DB plasmids, most containing a single MFD member among many.

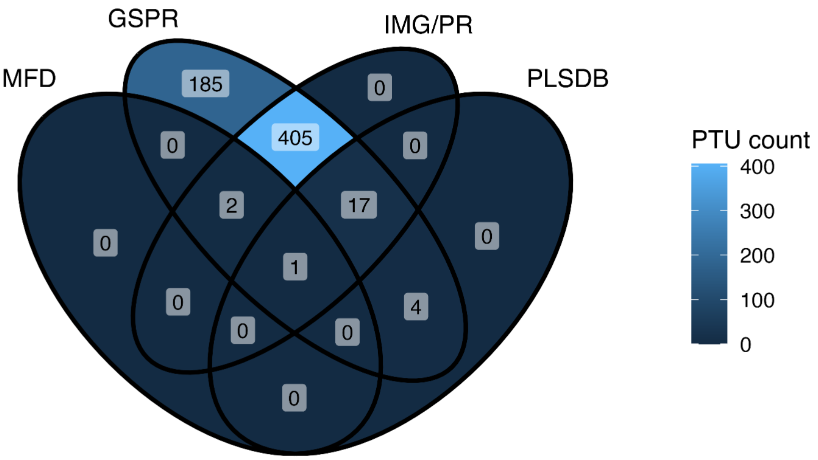


**Supplementary Figure 3.** Composition of PTUs containing plasmids from the Global Soil Plasmid Resource (GSPR) study. The analysis included the dereplicated and curated plasmid set of plasmids used throughout this study together with putatively complete plasmids from the GSPR study (v3.0) that were not already represented in our reference dataset. PTUs were inferred on this dataset using the same workflow described in this study **(Methods).** The Venn diagram shows only PTUs containing a GSPR plasmid. We investigated whether plasmids from the GSPR went into PTUs with plasmids from the MFD plasmidome, but found no direct overlap not already in our reference collection (IMG/PR and PLSDB), reflected by the absence of PTUs shared exclusively between MFD and GSPR plasmids in the Venn diagram.

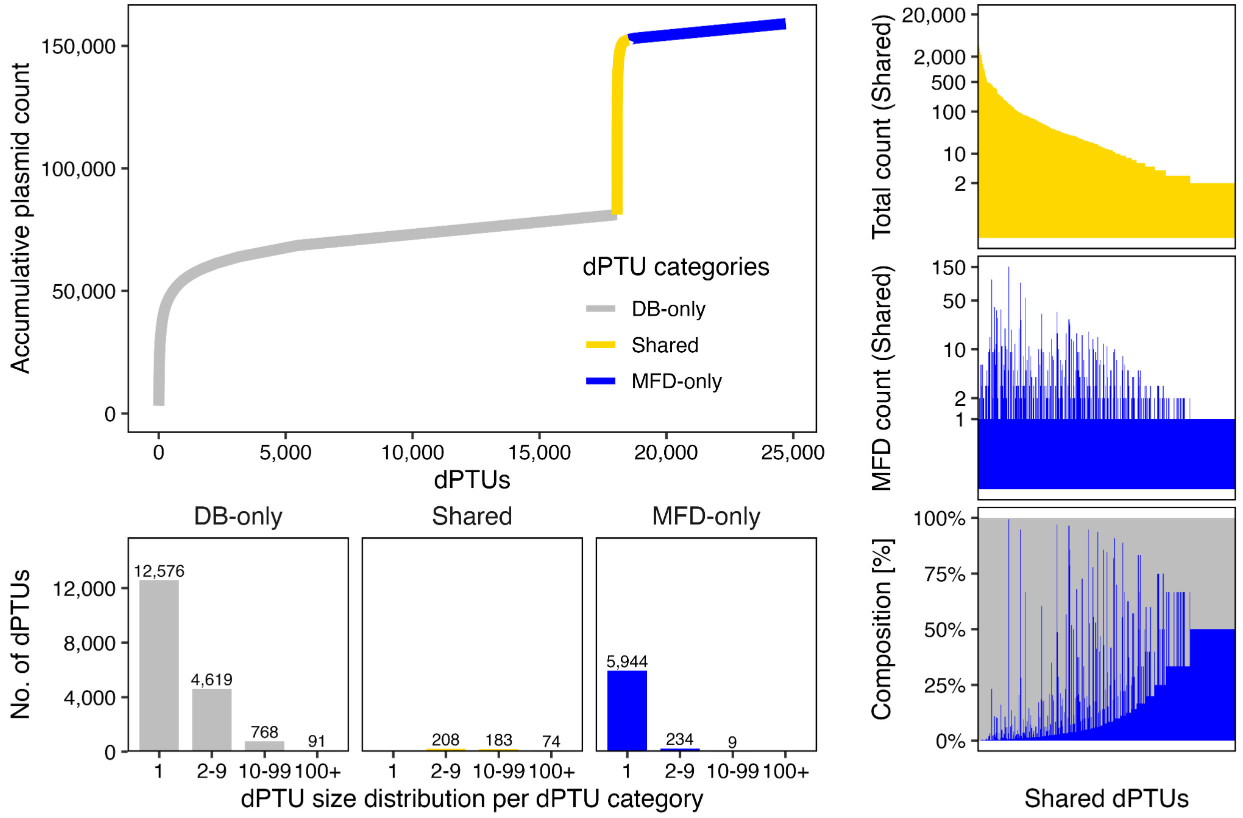


**Supplementary Figure 4.** Overview of the dPTU inference of MFD and reference collection plasmids. The 8,744 MFD and 150,308 DB plasmids were clustered into 24,705 dPTUs, each classified as DB-only), MFD-only, or Shared if a dPTU contained plasmids from both sources. The accumulation curve (top left) plots the cumulative number of plasmids as dPTUs are added in order of decreasing size. The bar plots (bottom left) give the number of MFD-only, DB-only and Shared PTUs within each size category. The three panels on the right give an overview of Shared PTUs, ordered from largest to smallest, showing the total count of plasmids per Shared PTU (top), the count of MFD plasmids in such Shared PTUs (middle) and the composition in percent of the number of plasmids from either source (bottom). Shared dPTUs were dominated by the reference plasmids, although more contained MFD plasmids (or multiple) than Shared PTUs **(Supplementary Fig. 2),** with some forming considerably larger groups. As dPTUs are based on amino acid similarity, these groups reflect MFD plasmids that are more similar in protein composition to the reference collection than is captured by the PTUs.

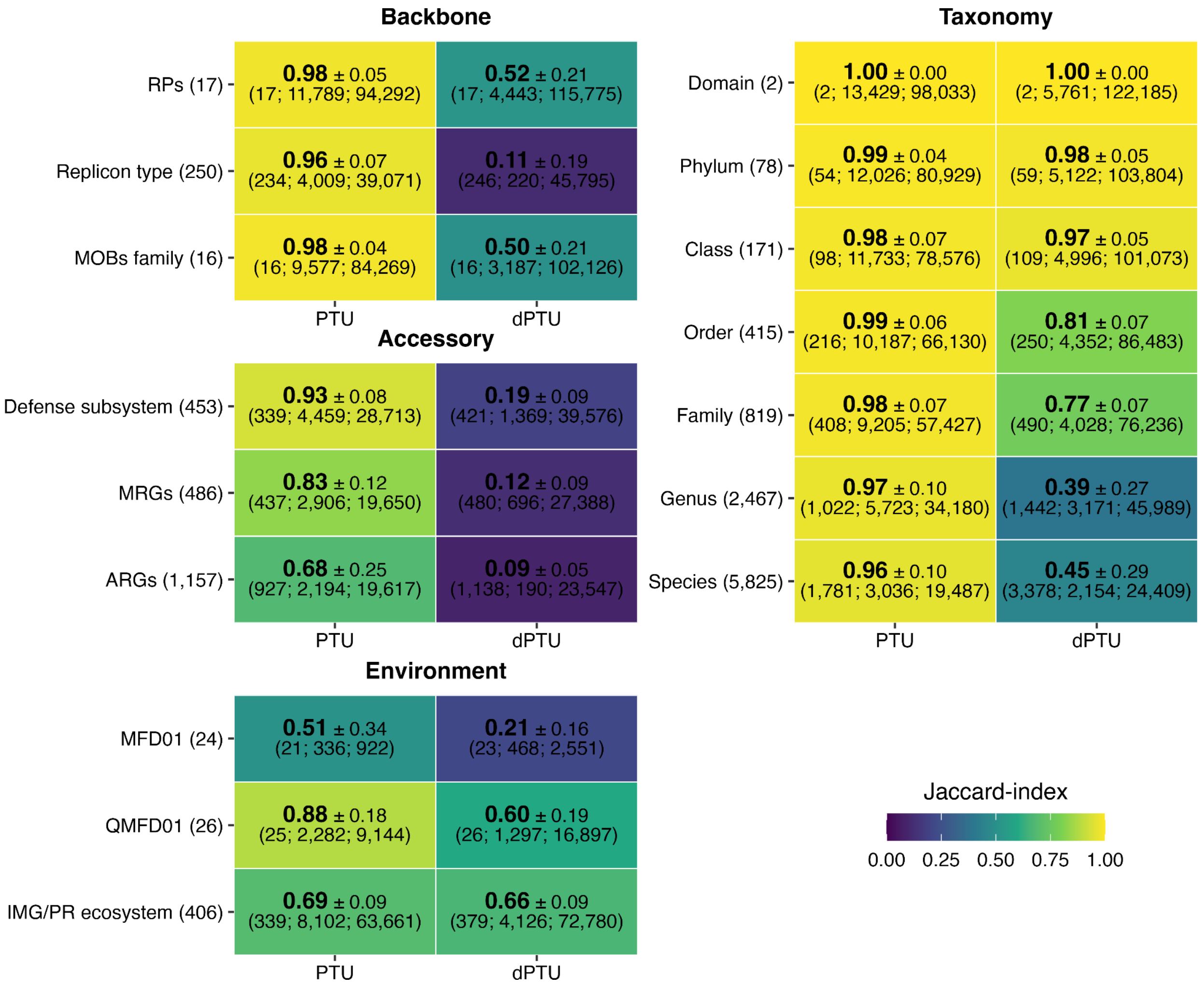


**Supplementary Fig 5.** Average similarity of plasmid traits across PTUs and dPTUs, measured using the Jaccard similarity. For each PTU or dPTU, a Jaccard similarity was calculated for the indicated trait, reflecting the extent to which members of the same group (PTU or dPTU) shared that trait. The values shown in the heatmap tiles represent the mean Jaccard value across groups, weighted by the number of members in each group; thus, larger groups contribute more strongly to the summary. Singletons were excluded. Higher values indicate greater consistency for the corresponding trait. The number in parentheses beside each trait label indicates the total number of categories within that trait. The three values in parentheses below each Jaccard similarity denote, in order, the number of those traits actually represented in groups with more than two members and without missing (NA) values, the number of such groups, and the number of plasmids represented in those groups.

**A**

| | MFD | IMG/PR | PLSDB |
|---|---|---|---|
| No. of plasmids | 8,744 | 100,439 | 49,869 |
| Domain | 8,712 (99.6%) | 85,931 (85.6%) | 45,377 (91.0%) |
| Phylum | 7,333 (83.9%) | 70,235 (69.9%) | 41,952 (84.1%) |
| Class | 6,835 (78.2%) | 68,340 (68.0%) | 40,970 (82.2%) |
| Order | 4,987 (57.0%) | 57,592 (57.3%) | 36,498 (73.2%) |
| Family | 4,188 (47.9%) | 49,799 (49.6%) | 33,539 (67.3%) |
| Genus | 2,221 (25.4%) | 35,095 (34.9%) | 16,687 (33.5%) |
| Species | 819 (9.4%) | 21,427 (21.3%) | 7,198 (14.4%) |

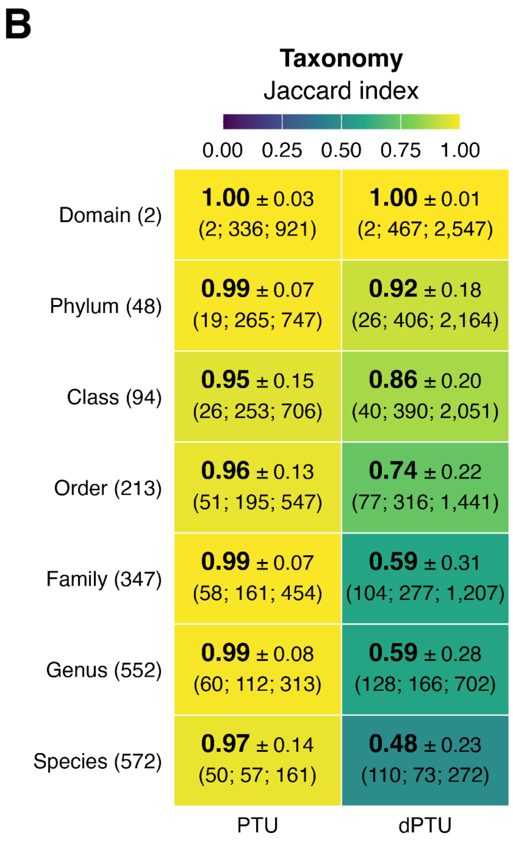


**Supplementary Fig. 6. (A)** Number of plasmids assigned at each taxonomic rank in MFD, IMG/PR and PLSDB. Taxonomy was assigned using MMseqs2 taxonomy with GTDB references. The top row gives the total plasmid count for each database; the remaining rows give the number and percentage assigned at each rank, from domain to species. **(B)** Weighted average Jaccard similarity ± weighted standard deviation of inferred host taxonomy within PTUs and dPTUs, both weighted by the number of MFD members included in each group, as described in Methods ("Biological coherence of PTUs and dPTUs"). Numbers in parentheses are explained in **Supplementary Fig. 5**. Only groups containing at least two MFD plasmids with taxonomic assignments at the corresponding rank were included.

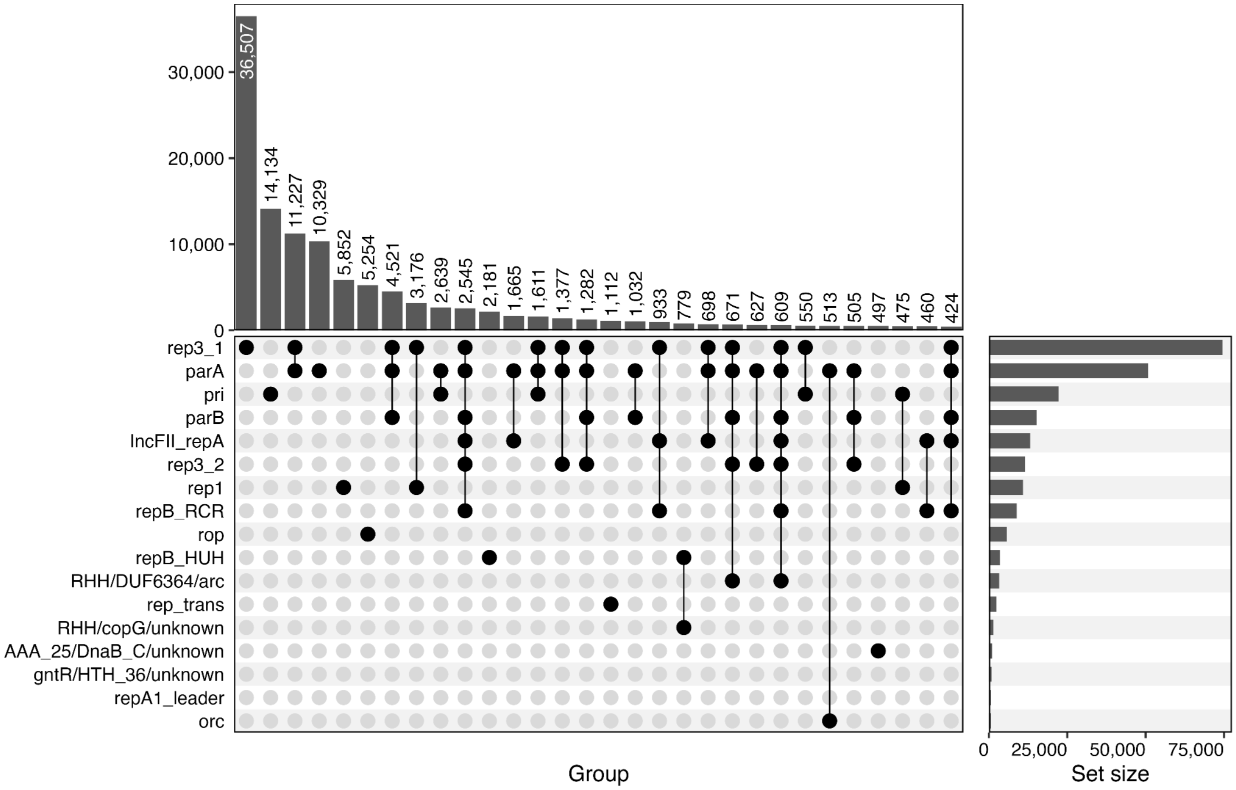


**Supplementary Figure 7.** UpSet plot displaying the co-occurrence of different replication-associated markers across the dereplicated plasmid set.

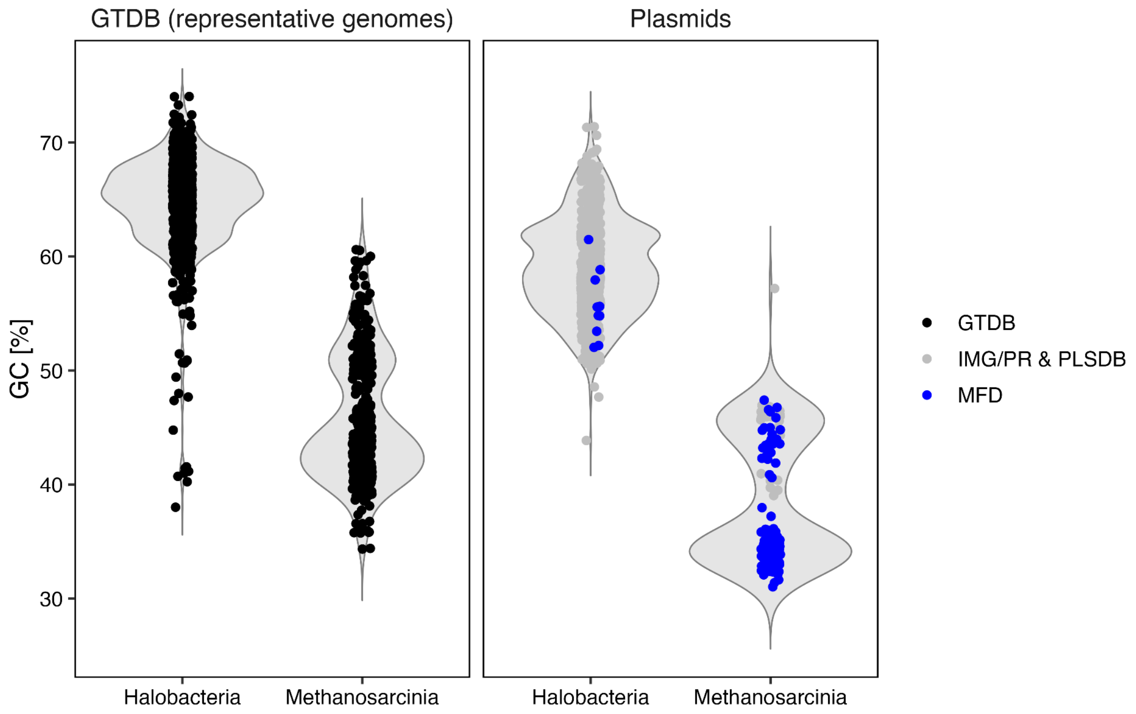


**Supplementary Fig. 8.** GC content of Halobacteria and Methanosarcinia representative genomes and plasmids. Violin plots show the distribution of GC content (%) for representative genomes from GTDB release 220 (left) and plasmids assigned to each class (right). Each point represents one genome or plasmid. Black points indicate GTDB genomes, grey points indicate IMG/PR and PLSDB plasmids, and blue points indicate MFD plasmids.

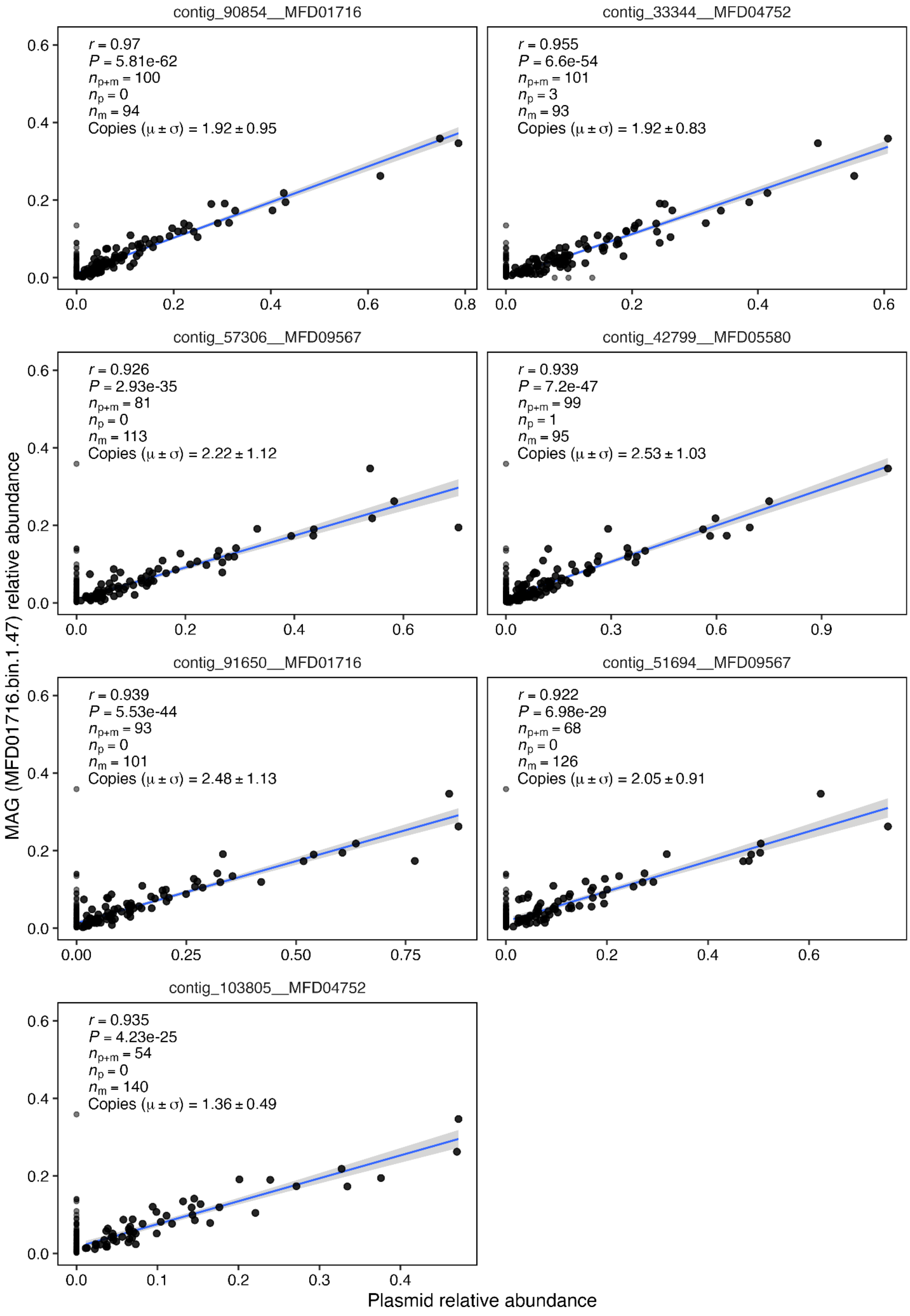
contig_90854__MFD01716
r = 0.97
P = 5.81e-62
n p+m = 100
n p = 0
n m = 94
Copies (μ ± σ) = 1.92 ± 0.95
contig_33344__MFD04752
r = 0.955
P = 6.6e-54
n p+m = 101
n p = 3
n m = 93
Copies (μ ± σ) = 1.92 ± 0.83
contig_57306__MFD09567
r = 0.926
P = 2.93e-35
n p+m = 81
n p = 0
n m = 113
Copies (μ ± σ) = 2.22 ± 1.12
contig_42799__MFD05580
r = 0.939
P = 7.2e-47
n p+m = 99
n p = 1
n m = 95
Copies (μ ± σ) = 2.53 ± 1.03
contig_91650__MFD01716
r = 0.939
P = 5.53e-44
n p+m = 93
n p = 0
n m = 101
Copies (μ ± σ) = 2.48 ± 1.13
contig_51694__MFD09567
r = 0.922
P = 6.98e-29
n p+m = 68
n p = 0
n m = 126
Copies (μ ± σ) = 2.05 ± 0.91
contig_103805__MFD04752
r = 0.935
P = 4.23e-25
n p+m = 54
n p = 0
n m = 140
Copies (μ ± σ) = 1.36 ± 0.49
MAG (MFD01716.bin.1.47) relative abundance
Plasmid relative abundance

85

**Supplementary Fig. 9.** Co-abundance of the putative Methanosarcina plasmids and the Methanosarcina MAG MFD01716.bin.1.47. Each panel shows one plasmid (panel title), plotting the relative abundance of the plasmid (x-axis) against the relative abundance of the MAG (y-axis), estimated with sylph. Each point is one MFD metagenome in which the plasmid, the MAG or both were detected. The blue line is a linear regression with its 95% confidence interval (grey), fitted only to samples where both the plasmid and the MAG were detected. In the inset table, *r* is the Pearson correlation coefficient between plasmid and MAG relative abundance across samples where both were detected, and P is the corresponding two-sided P value. The sample counts are $n_{p+m}$ for samples where both were detected, $n_p$ for samples where only the plasmid was detected, and $n_m$ for samples where only the MAG was detected. Copies (μ ± σ) is the mean ± standard deviation of the per-sample plasmid to MAG relative abundance ratio over the $n_{p+m}$ detected samples.

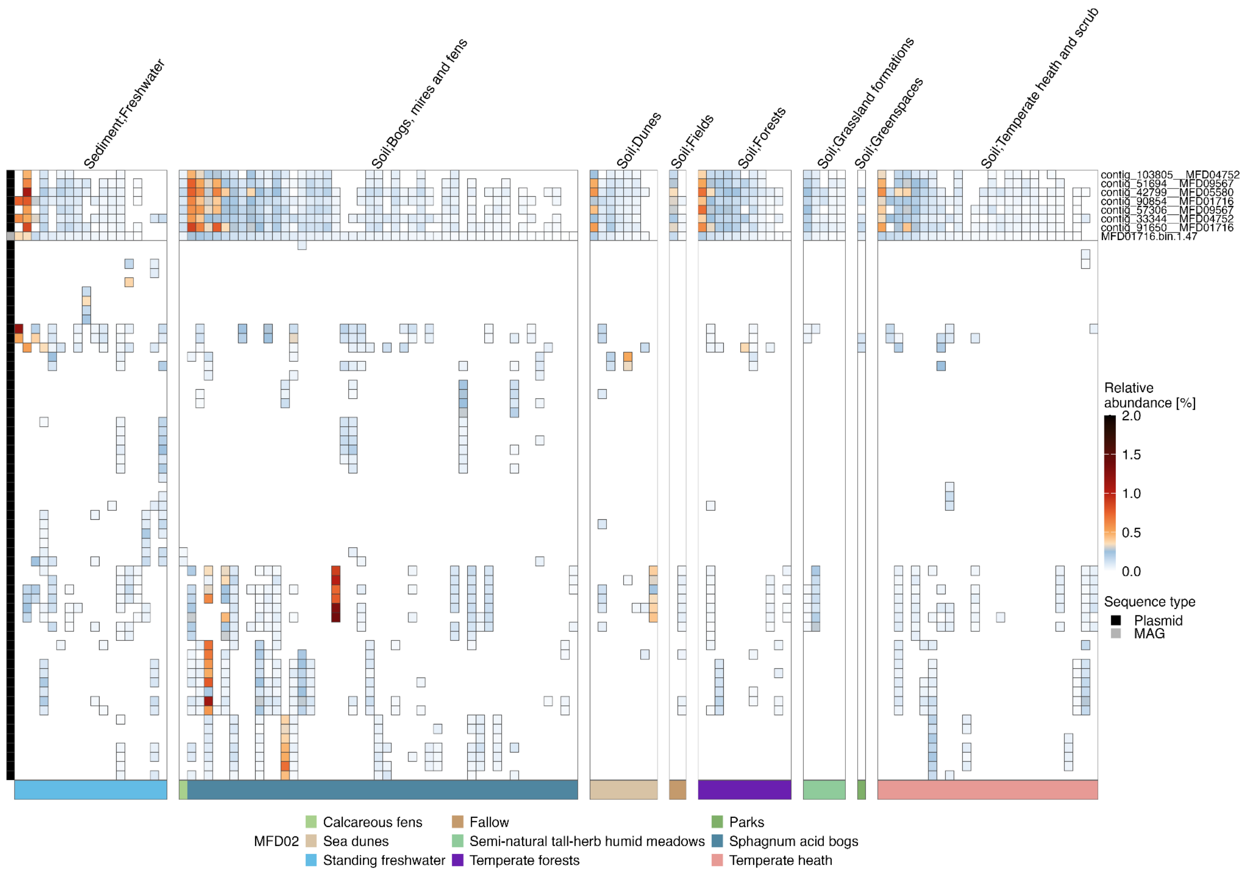


**Supplementary Figure 10.** Relative abundance of *Methanosarcina* plasmids and its co-occurring plasmids across Danish terrestrial metagenomes. The heatmap depicts the sylph-estimated relative abundance (%) for a metagenome-assembled genome (MAG; MFD01716.bin.1.47) and associated plasmid contigs across Microflora Danica (MFD) metagenomes. Each column is a sample and each row a plasmid (except the highlighted MAG); tile colour denotes relative abundance, scaled from white (low) through blue and orange to black. The top block highlights the MAG and its seven most closely associated plasmid contigs (row labels shown); the lower block shows the remaining plasmids. The heatmaps are filtered to show the putative *Methanosacina* plasmids **(Fig 4)** in samples that have ≥5 detected plasmids. The left annotation bar indicates element origin (MAG, red; plasmid, blue). Samples are grouped by sample type and primary habitat (MFD level-1 classification, panel headers) and, within each group; the coloured bar beneath the heatmap gives the finer habitat assignment (MFD level-2 classification, "MFD02"). Rows are ordered by hierarchical clustering (Ward's method on Bray-Curtis distances of Hellinger-transformed abundances).

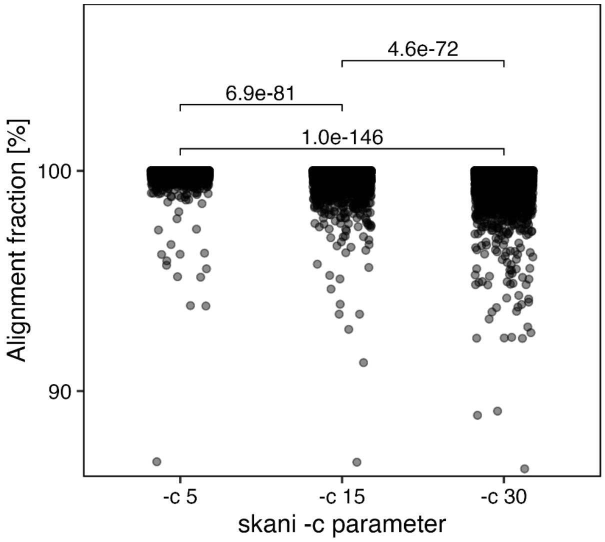


**Supplementary Fig. 11:** Alignment fractions for the curated set of 8,744 putatively complete MFD plasmids, calculated using skani (v0.2.2; dist -m 50 -s 70) with -c 5, -c 15, and -c 30. skani triangle --diagonal was used for PTU inference (see section “PTU inference”), where diagonal (self-to-self) alignment fractions are set to 100% by default. Here, dist was used to calculate the diagonal alignment fractions, where each plasmid is compared with itself and an alignment fraction of 100% is expected (but not default). The average alignment fractions were 99.93%, 99.87%, and 99.78%, respectively. The differences were small, but -c 5 produced an average significantly closer to 100% than -c 15 or -c 30 (two-sided paired t-tests with Bonferroni correction). This was used as justification to use -c 5 for the PTU inference to improve the accuracy.

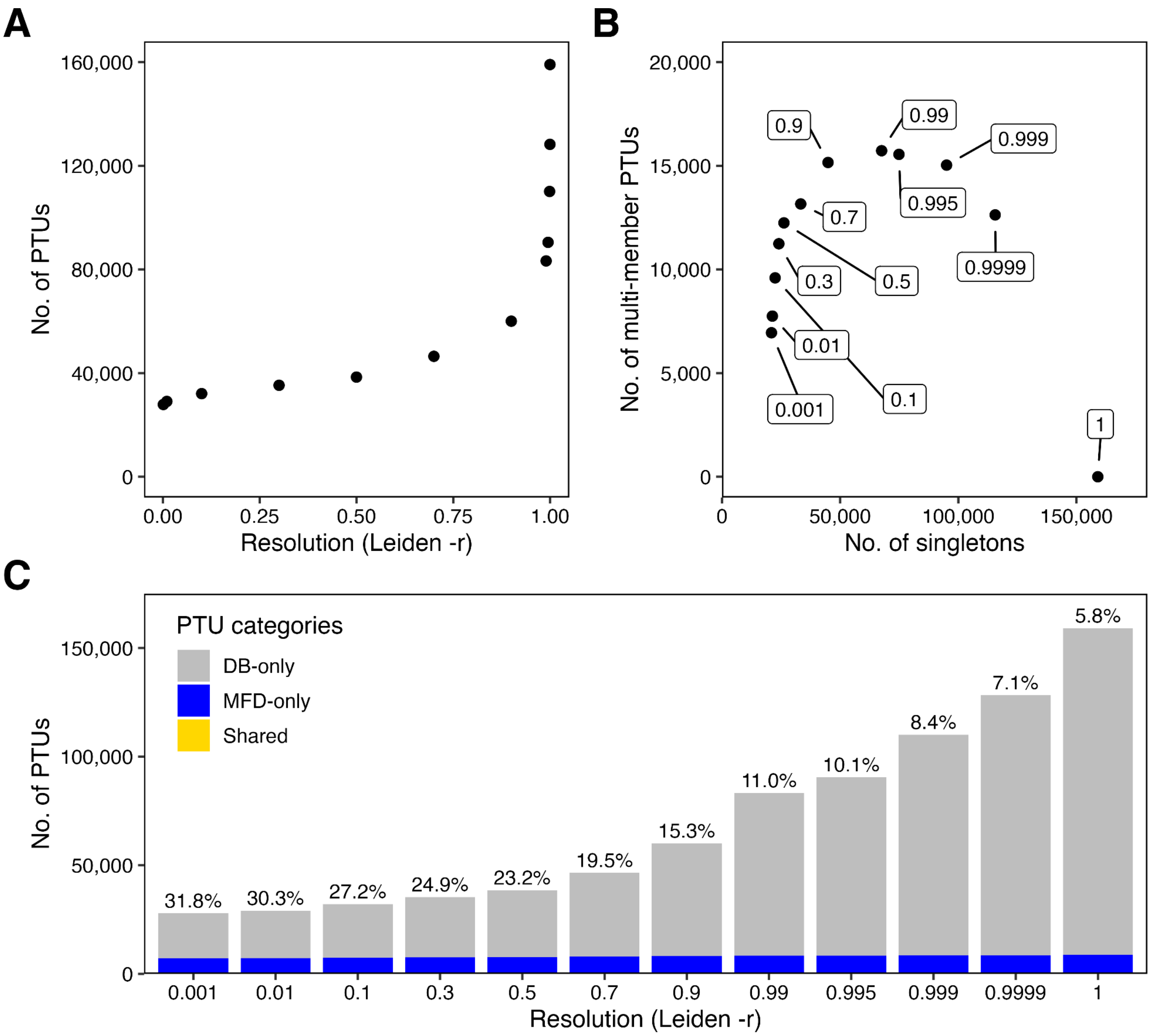


**Supplementary Fig. 12.** Selection of the Leiden resolution parameter for plasmid taxonomic unit (PTU) inference. Effect of the Leiden resolution parameter (-r) on the plasmid network partition, used to justify the chosen value of 0.9. **A)** Total number of PTUs as a function of resolution; group counts rise slowly at low resolution and increase sharply as -r approaches 1. **B)** Number of multi-member plasmid clusters (≥2 members) versus number of singletons across the same resolution sweep (each point labelled with its -r value). Beyond -r = 0.9, increasing resolution reduces the number of multi-member PTUs while the number of singletons continues to increase, and at -r = 1 almost all elements are singletons (i.e. added resolution fragments existing PTUs rather than resolving new ones). **C)** PTU counts by resolution, coloured by source; DB-only (grey), MFD-only (blue) and Shared (yellow). Percentages give the novelty rate (MFD-only PTUs as a fraction of the total). Across all resolutions, the Shared fraction remains negligible, showing that MFD plasmids stay largely unique to MFD regardless of partition granularity.
