## Supplementary Notes 1-2 for "Nation-scale recovery of complete environmental plasmids pushes the boundaries of plasmid diversity through long-read sequencing"

Bertil Sørensen[1^], Connor L. Brown[1^], Mantas Sereika[1], Søren Heidenbach[1], Philip Hugenholtz[1,2], Mads Albertsen[1], Francesco Delogu[1,3]*
[1]Center for Microbial Communities, Aalborg University, Denmark
[2]Australian Centre for Ecogenomics, School of Chemistry and Molecular Biosciences, The University of Queensland, Brisbane, Queensland, Australia
[3]Department of Computer Science, Aalborg University, Aalborg, Denmark
^Shared first authorship
*Corresponding author

#### Supplementary Note 1

**Replication-associated protein family curation**

Many plasmids use a winged helix-turn-helix replication initiation protein (RP) in part exemplified by the Rep3_N/Rep3_C[1], however other replication initiation mechanisms exist, and replication itself involves coordinated effort of many host- or plasmid-derived proteins. To propose new hallmark genes for annotating plasmids, and to characterize replicon diversity of our plasmid set, we devised a scheme to recover major RP families through the combination of the Nishimura *et al.*[1] protein set and mobileOG-db **(Supplementary Figure X1).** Briefly, a pre-release version of mobileOG-db v.2 (clustered at 90% identity) was analyzed to recover putative RPs. Sequences from mobileOG-db were combined with the Nishimura *et al.*[1] sequences and then the merged set were compared to the MFD and DB plasmid proteins using diamond blastp (minimum identity 40% and query-cover of 50%). The intermediate potential RP set encompassing mobileOG-db sequences, Nishimura *et al.*[1], and MFD/DB plasmid sequences were then clustered with SwissProt using mmseqs (--min-seq-id 0.4 -c 0.5 --cov-mode 0). This produced a list of 1,074 SwissProt entries associated to the potential RP set. These entries were manually inspected to produce a list of 100 SwissProt entries potentially associated with plasmid replication which in turn were associated with 162,925 potential RPs. We then used an iterative refinement strategy where sequences were clustered and the clusters merged if any detectable sequence homology was found (via diamond blastp --ultra-sensitive) between cluster representatives. Within each of the final 17 clusters, the Levenstein distance of each member was calculated relative to the cluster representative and members with distance greater than 4 standard deviations than the cluster mean were excluded.

We next characterized the domain attributes **(Supplementary Table X1)** and matches to reference sequences of the protein families **(Supplementary Table X2)** and performed an annotation criteria parameter sweep to estimate the number of potential RPs encoded across plasmids **(Supplementary Figures X1-X3)**. Each protein family was dominated by a characteristic PFAM domain: parA by PF09140 (ParA/MinD-type ATPase, 6,502 hits); rep3_1 and rep3_2 both by PF01051 (Rep_3 replication initiator); rep1 by PF01446 (PriCT-1 / Rep_1, 1,247 hits); and orc by PF00270 (the DEAD-box helicase-associated ORC domain, 352 hits). The impact of minimum percent identity and query cover when using the RP set was variable across protein families in returned hits, with short leader peptides (e.g., rop and repA_1_leader) having no loss at increasingly greater levels of stringency. However, these proteins represent a small subset of the RPs, and this result likely reflects the sequence-specificity required for the peptide to perform its function. In sum, these results show the relative divergence within each specific detected RP and should be a starting point for using the RP set for annotation of plasmids in the future.

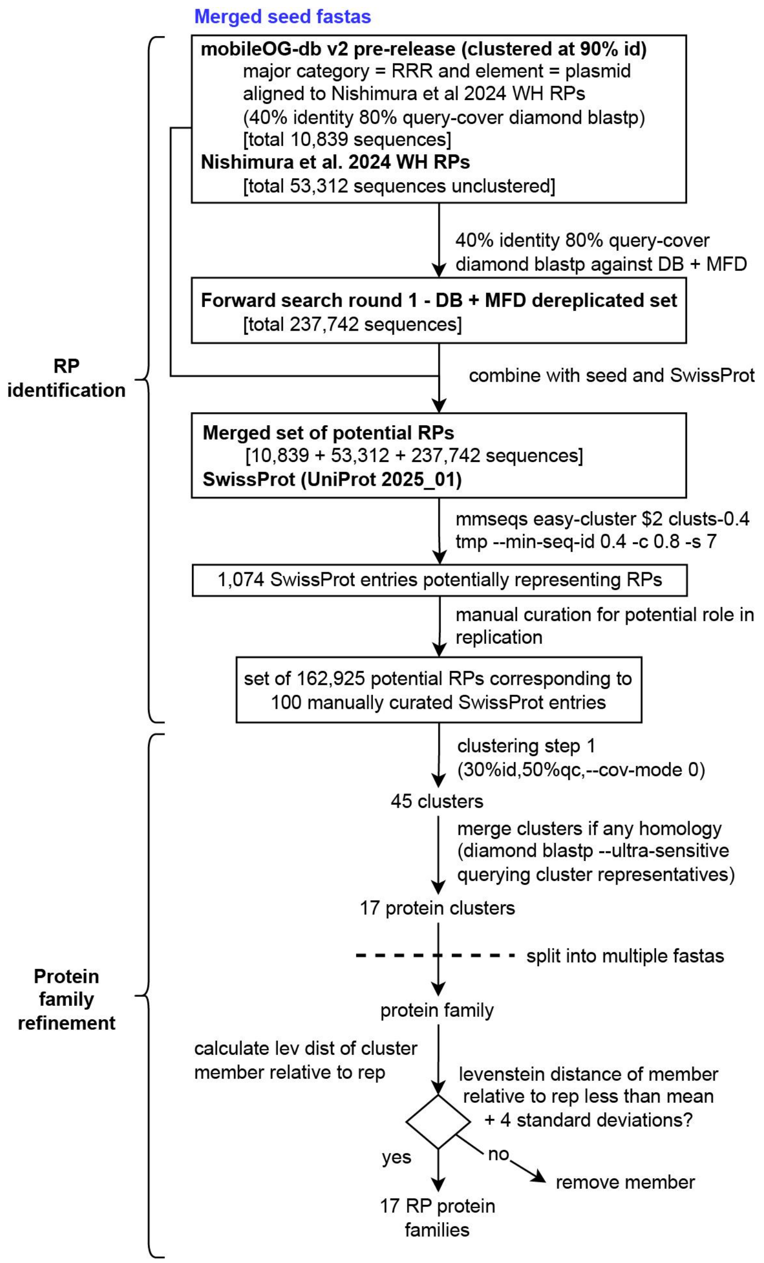


**Supplementary Figure X1.** Curation and refinement of 17 RP protein families.

**Supplementary Table X1. Annotations of 17 RP families and proposed gene names.**

| Representative sequence | Proposed gene name (current study) | Most common PFAM(s) |
|---|---|---|
| IMGPR_plasmid_3300000136_000001_7 | rep3_1 | Rep3_N |
| NZ_CP140809.1_1 | parA | AAA_31,ArsA_ATPase,CbiA,CBP_BcsQ,Fer4_NifH,MipZ,ParA |
| IMGPR_plasmid_3300053491_019932_57 | IncFII_repA | IncFII_repA |
| NZ_CP134741.1_1 | rep3_2 | Rep3_C,Rep3_N |
| NZ_KM877517.1_1 | repB_RCR | RepB-RCR_reg |
| IMGPR_plasmid_3300048591_000433_113 | parB | ParB_N |
| NZ_CP041342.1_1 | rop | Rop |
| mobileOG_002523916 | rep1 | Rep_1 |
| IMGPR_plasmid_2894567016_000001_12 | pri | PriCT_1 |
| IMGPR_plasmid_3300044557_000615_2 | rep_trans | RepD-like_N,Rep_trans |
| IMGPR_plasmid_3300053491_067451_1 | repB_HUH | RepB_C,Rep_OBD |
| IMGPR_plasmid_3300026519_000008_4 | AAA_25/DnaB_C/unknown | AAA_25,DnaB_C |
| NZ_CP106969.1_91 | orc | AAA,AAA_16,AAA_22,Cdc6_C,Cdc6_lid,NPHP3_N |
| IMGPR_plasmid_3300053491_067267_3 | RHH/copG/unknown | RHH_5 |
| IMGPR_plasmid_2913276826_000002_1 | gntR/HTH_36/unknown | GntR,HTH_36 |
| NZ_CP115017.1_39 | RHH/DUF6364/arc | Arc,DUF6364 |
| NZ_LR882059.1_90 | repA1_leader | RepA1_leader |

**Supplementary Table X2. Allocation of specific SwissProt annotations to RP gene families.**

| Best hit to SwissProt (RP current study) | Proposed gene name (current study) | Swiss Prot ID | Gene name (SP) | SwissProt annotation |
|---|---|---|---|---|
| NZ_OR237799.1_1 | IncFII_repA | P18023 | repZ | REPZ_ECOLX Replication initiation protein OS=Escherichia coli OX=562 GN=repZ PE=4 SV=1 |
| NZ_OX030730.1_1 | IncFII_repA | Q52221 | repA | REP7_ECOLX Replication initiation protein OS=Escherichia coli OX=562 GN=repA PE=1 SV=1 |
| OW967847.1_31 | IncFII_repA | P03066 | repA | REPA2_ECOLX Replication initiation protein OS=Escherichia coli OX=562 GN=repA PE=3 SV=1 |
| NZ_CP146626.1_201 | orc | Q9HHR1 | orc5 | CDC65_HALSA ORC1-type DNA replication protein 5 OS=Halobacterium salinarum (strain ATCC 700922 / JCM 11081 / NRC-1) OX=64091 GN=orc5 PE=3 SV=2 |
| CP057153.1_53 | parA | P07620 | parA | PARA_ECOLX Plasmid partition protein A OS=Escherichia coli OX=562 GN=parA PE=1 SV=1 |
| AP028980.1_7 | parB | P07621 | parB | PARB_ECOLX Plasmid Partition par B protein OS=Escherichia coli OX=562 GN=parB PE=3 SV=1 |
| NZ_OZ038807.1_188 | parB | P62559 | sopB | SOPB_ECO57 Protein SopB OS=Escherichia coli O157:H7 OX=83334 GN=sopB PE=3 SV=1 |
| mobileOG_000001406 | pri | P14752 | repS | REPS_STRPY Protein RepS OS=Streptococcus pyogenes OX=1314 GN=repS PE=4 SV=1 |
| mobileOG_000001160 | pri | P18016 | rep | REP_CLOPF Replication protein OS=Clostridium perfringens OX=1502 GN=rep PE=4 SV=1 |
| IMGPR_plasmid_2693429673_000001_2 | repD/Rep_trans | P14490 | repM | REPM_STAAU Replication initiation protein OS=Staphylococcus aureus OX=1280 GN=repM PE=3 SV=1 |
| IMGPR_plasmid_3300053491_005082_4 | repD/Rep_trans | P19529 | repN | REPN_STAAU Replication initiation protein OS=Staphylococcus aureus OX=1280 GN=repN PE=1 SV=1 |
| NZ_CP128502.1_3 | rep1 | P13963 | rep | REP_BACAM Protein rep OS=Bacillus amyloliquefaciens OX=1390 GN=rep PE=3 SV=1 |
| mobileOG_003363156 | rep3_1 | P05684 | repC | REPC_RHIRH Putative replication protein C OS=Rhizobium rhizogenes OX=359 GN=repC PE=4 SV=1 |
| NZ_OQ683451.1_9 | rep3_1 | P07676 | tfrA | TRFA_ECOLX Plasmid replication initiator protein TrfA OS=Escherichia coli OX=562 GN=trfA PE=1 SV=1 |

| | | | | |
|---|---|---|---|---|
| mobileOG_000324082 | rep3_1 | A0A011QFS8 | AW11_02368 | A0A011QFS8_ACCRE Replication initiator protein A OS=Accumulibacter regalis OX=522306 GN=AW11_02368 PE=4 SV=1 |
| IMGPR_plasmid_640048375_000001_2 | rep3_1 | P17212 | NA | 41KD_LACHE 41 kDa protein OS=Lactobacillus helveticus OX=1587 PE=4 SV=1 |
| mobileOG_000003710 | rep3_1 | P17492 | NA | REPA_NEIGO Replication protein OS=Neisseria gonorrhoeae OX=485 PE=3 SV=1 |
| IMGPR_plasmid_651053003_000002_5 | rep3_1 | P10022 | NA | YGI1_BACTU Uncharacterized 23.6 kDa protein OS=Bacillus thuringiensis OX=1428 PE=4 SV=2 |
| CP139865.1_4 | rep3_1 | P13968 | repL | REPLM_BACIU Replication and maintenance protein OS=Bacillus subtilis OX=1423 GN=repL PE=4 SV=1 |
| NZ_PP215916.1_61 | rep3_2 | P0A254 | repA | REP12_ECOLX RepFIB replication protein A OS=Escherichia coli OX=562 GN=repA PE=3 SV=1 |
| IMGPR_plasmid_3300053491_058467_2 | repB_HUH | P20045 | repB | REPB_LACPN Replication protein RepB OS=Lactiplantibacillus plantarum OX=1590 GN=repB PE=3 SV=1 |
| OZ026790.1_85 | repB_RCR | P03847 | repA2 | COPB2_ECOLX Replication regulatory protein repA2 OS=Escherichia coli OX=562 GN=repA2 PE=4 SV=2 |
| NZ_PP261916.1_104 | RHH/DUF6364/arc | P10030 | pifC | PIFC_ECOLI Transcriptional repressor PifC OS=Escherichia coli (strain K12) OX=83333 GN=pifC PE=1 SV=1 |
| k127_77639__MFD10529_9 | AAA_25/DnaB_C/unknown | P20356 | repA | REPJ_ECOLX Regulatory protein RepA OS=Escherichia coli OX=562 GN=repA PE=1 SV=1 |
| IMGPR_plasmid_2913276826_000001_1 | gntR/HTH_36/unknown | P15394 | repA | REPA_AGRTU Replicating protein OS=Agrobacterium tumefaciens OX=358 GN=repA PE=4 SV=1 |
| NZ_MZ396393.1_2 | RHH/copG/unknown | P13920 | copG | COPG_STRAG Protein CopG OS=Streptococcus agalactiae OX=1311 GN=copG PE=1 SV=1 |
| NZ_CP023850.1_1 | repA1_leader | P62550 | repL | REPL2_ECOLX Positive regulator of RepFIC repA1 expression OS=Escherichia coli OX=562 GN=repL PE=4 SV=1 |

52

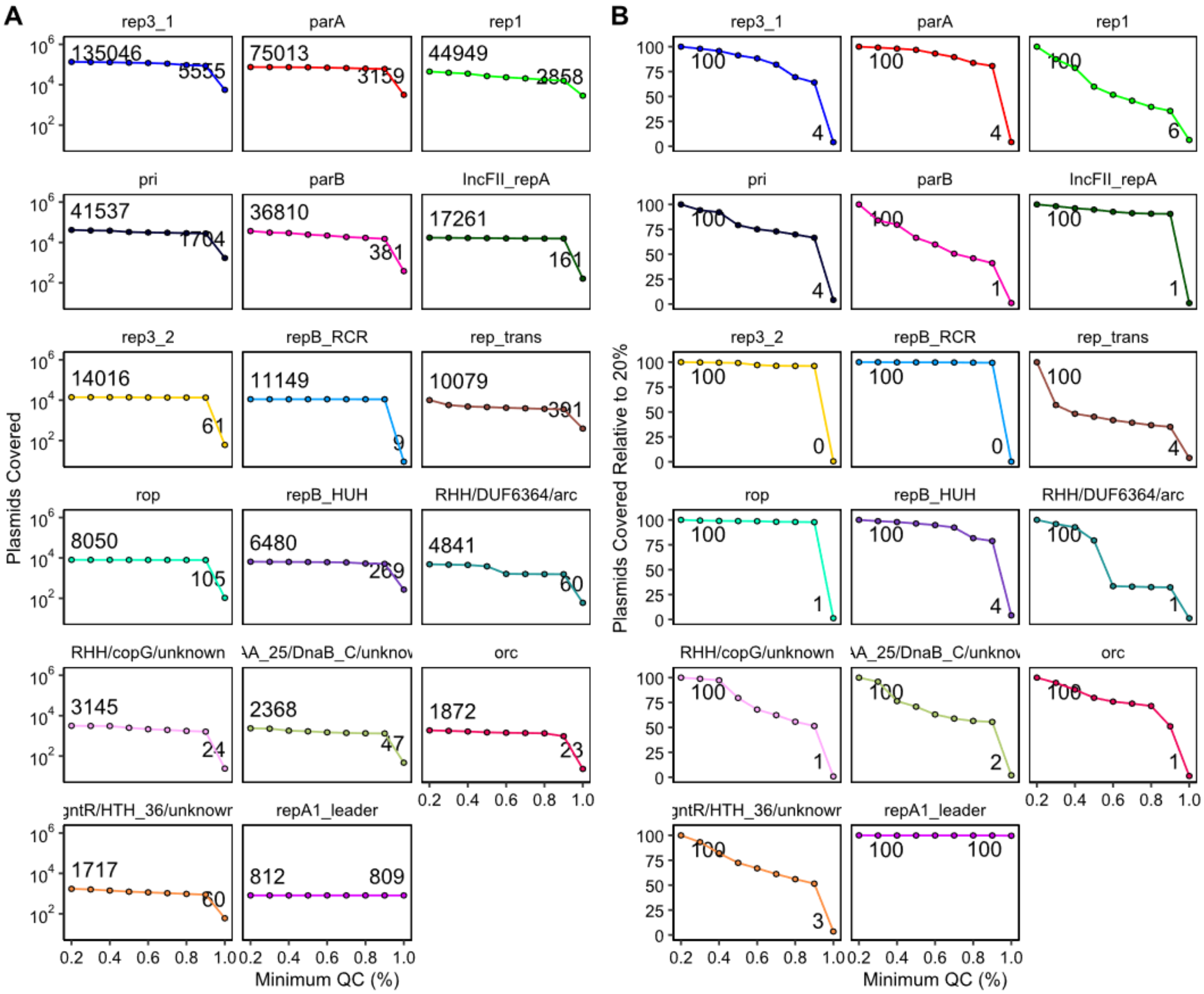


**Supplementary Figure X2. Query cover (QC) parameter sweep partitioned by RP family. (A)** Cumulative counts of plasmids from MFD + DB encompassed by the RP family across different QC values. **(B)** Percentage of plasmids encompassed by the RP family at a given QC value relative to the minimum (0.2).

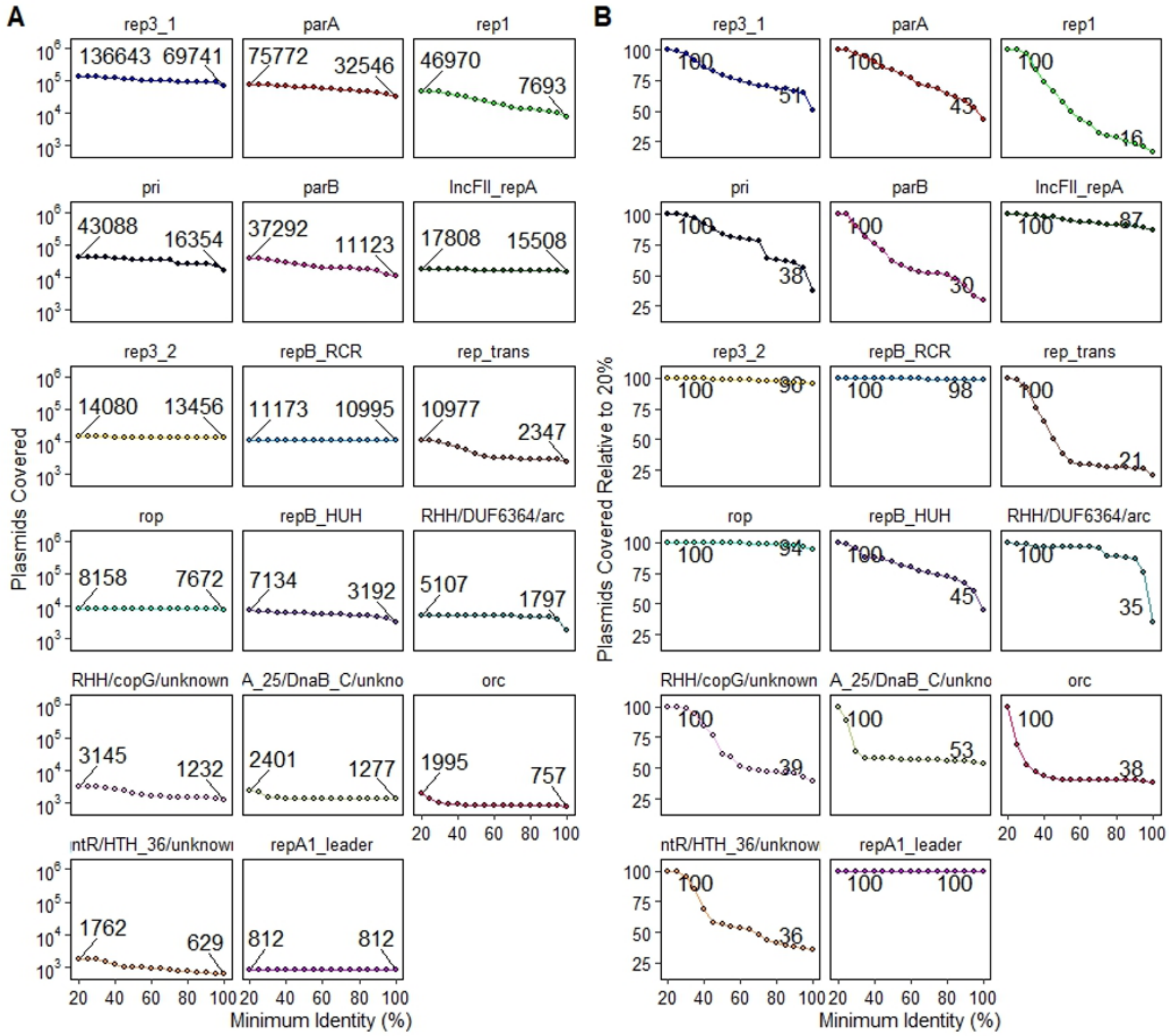


**Supplementary Figure X2. Alignment identity parameter sweep partitioned by RP family.** **(A)** Cumulative counts of plasmids from MFD + DB encompassed by the RP family across different minimum identities. **(B)** Percentage of plasmids encompassed by the RP family at a given minimum identity value relative to those encompassed by minimum identity 20%.

Range (Min, Max, Median) in Copy Number

| | 20 % min | | 40 % min | | 80 % min | | 90 % min | |
|---|---|---|---|---|---|---|---|---|
| | all plasmids | just MFD | all plasmids | just MFD | all plasmids | just MFD | all plasmids | just MFD |
| AAA_25/DnaB_C/unknown | 1,8,1 | 1,2,1 | 1,8,2 | 1,2,1 | 1,8,2 | 1,2,1 | 1,8,2 | 1,2,1 |
| rop | 1,11,1 | 1,1,1 | 1,11,1 | 1,1,1 | 1,11,1 | 1,1,1 | 1,11,1 | 1,1,1 |
| RHH/DUF6364/arc | 1,5,1 | 2,2,2 | 1,5,1 | | 1,5,1 | | 1,5,1 | |
| RHH/copG/unknown | 1,4,1 | 1,3,1 | 1,4,1 | 1,2,1 | 1,4,1 | 1,2,1 | 1,4,1 | 1,2,1 |
| repB_RCR | 1,3,1 | 1,1,1 | 1,3,1 | 1,1,1 | 1,3,1 | 1,1,1 | 1,3,1 | 1,1,1 |
| repB_HUH | 1,6,1 | 1,2,1 | 1,6,1 | 1,2,1 | 1,6,1 | 1,1,1 | 1,6,1 | 1,1,1 |
| repA1_leader | 1,1,1 | | 1,1,1 | | 1,1,1 | | 1,1,1 | |
| rep3_2 | 1,7,1 | 1,2,1 | 1,7,1 | 1,2,1 | 1,7,1 | 1,2,1 | 1,7,1 | 1,2,1 |
| rep3_1 | 1,75,1 | 1,34,1 | 1,75,1 | 1,16,1 | 1,50,1 | 1,6,1 | 1,42,1 | 1,6,1 |
| rep1 | 1,14,1 | 1,5,1 | 1,14,1 | 1,5,1 | 1,14,1 | 1,3,1 | 1,14,1 | 1,3,1 |
| rep_trans | 1,7,1 | 1,2,1 | 1,6,1 | 1,1,1 | 1,6,1 | 1,1,1 | 1,5,1 | 1,1,1 |
| pri | 1,35,1 | 1,7,1 | 1,35,1 | 1,7,1 | 1,18,1 | 1,7,1 | 1,18,1 | 1,7,1 |
| parB | 1,34,1 | 1,4,1 | 1,31,1 | 1,3,1 | 1,8,1 | 1,2,1 | 1,8,1 | 1,1,1 |
| parA | 1,49,1 | 1,18,1 | 1,28,1 | 1,17,1 | 1,27,1 | 1,11,1 | 1,25,1 | 1,9,1 |
| orc | 1,15,1 | 1,5,1 | 1,15,1 | 1,5,1 | 1,11,1 | 1,5,1 | 1,10,1 | 1,5,1 |
| IncFII_repA | 1,7,1 | 1,2,1 | 1,7,1 | 1,2,1 | 1,7,1 | 1,2,1 | 1,6,1 | 1,2,1 |
| gntR/HTH_36/unknown | 1,8,1 | 1,8,1 | 1,8,1 | 1,8,1 | 1,2,1 | 1,2,1 | 1,2,1 | 1,2,1 |

**Supplementary Figure X3.** Effect of minimum percent identity relative to reference for the plasmid markers

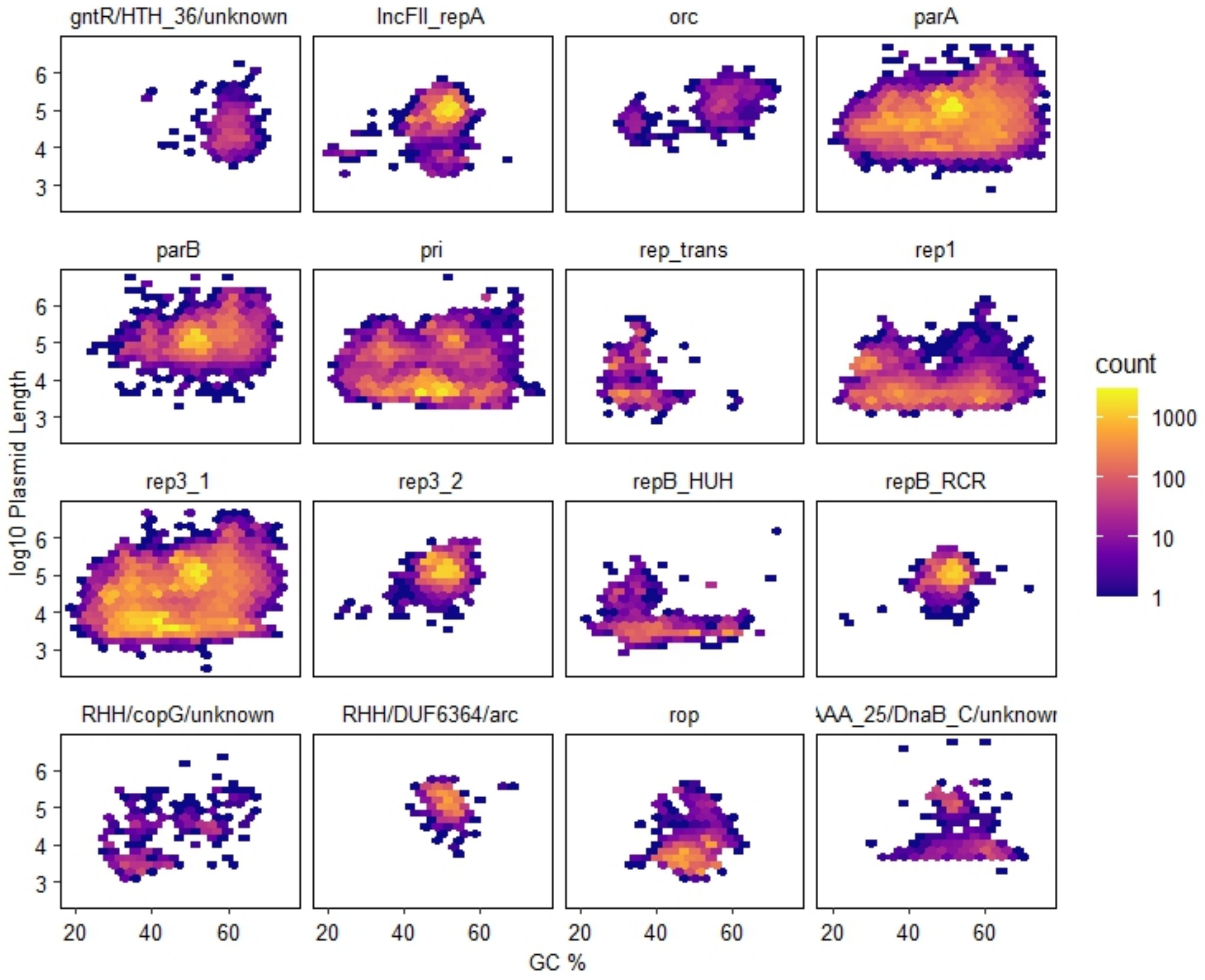


**Supplementary Figure X4.** Sequence length and GC% content characteristics of plasmids from various RP families.

**Functional annotation of major RP families**

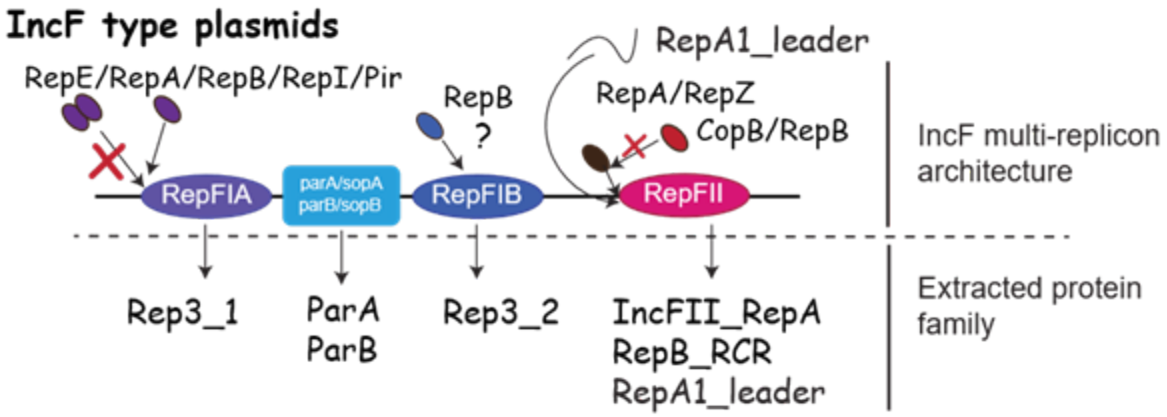


**IncF-associated replication proteins**

IncF-type plasmids variably encode some combination of three different replicons, RepFIA, RepFIB, and RepFII[2]. Each replicon is associated with a different set of RPs. Replication initiation at RepFIA, which can be found on the IncF-plasmid F, typified by RepE (P03856[3]) is initiated by monomeric winged helix-turn-helix (wHTH) protein and deregulated by the dimeric form. Less is known about the initiation mechanism of RepFIB[2], but its presence is sufficient to sustain replication in the absence of RepFIA. The last replicon, IncFII, is controlled by the activity of a negative regulator, CopB/RepB (here termed RepB_RCR) typified by P03847[4]. In plasmid F, the replicon is inactivated by transposon insertion but in related plasmid R1, it is the only active replicon[5]. It requires the activity of IncFII_RepA (P03066[6] on plasmid R100) for replication[7]. In addition, IncF and related plasmids often encode partitioning proteins ParA and ParB (also known as SopA/SopB). These proteins mediate localization of the plasmid to the poles of the cell during cell division, ensuring distribution of plasmid copies to daughter cells[8].

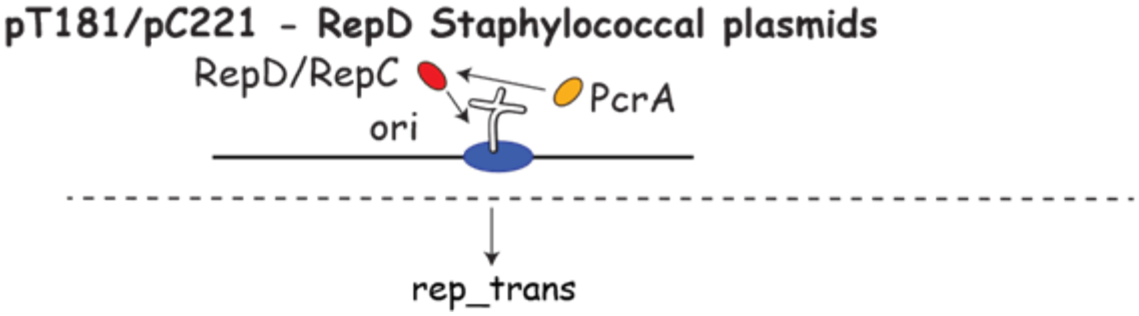


**Alternative replication initiator RepD**

Small Staphylococcal plasmids such as pT181/pC221 utilize an unusual replication initiator protein RepD (pT181) or RepC (pC221)[9]. A dimeric version of the protein binds to a sequence specific double stranded origin site and forms a transient cruciform structure, revealing its single-stranded nick site that is acted upon by the rep protein. This then aids recruitment of helicase PcrA. The rep_trans name is proposed based on the most common PFAM of this protein class.

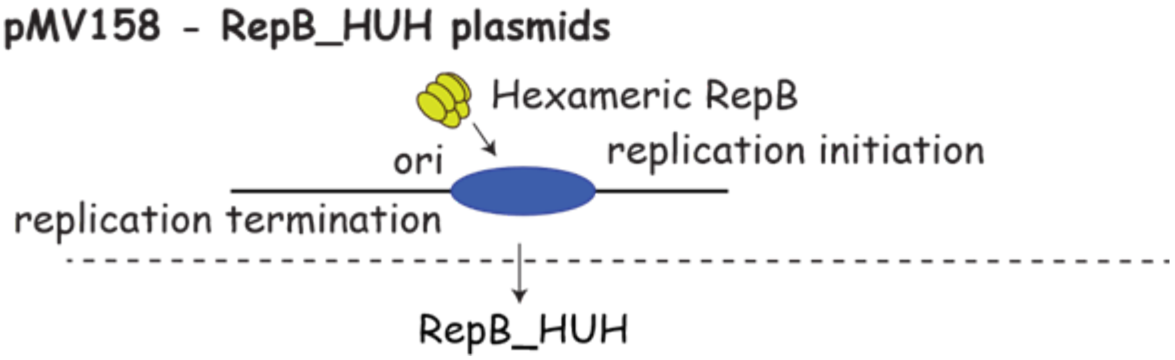


**Alternative replication initiator RepB_HUH**

Streptococcal plasmid pMV158 uses a distinct replication initiation protein, RepB, which is distinct for its hexameric configuration and use of a recombinase HUH motif for its activity. It is also distinct in that it catalyzes both the replication initiation and the termination step[10].

**Other proteins**

**Archaeal plasmid replication initiator orc**

Archaeal plasmids replicate using distinct machinery which, in part, resembles the basic machinery of linear chromosome replication of eukaryotes[11]. The primary replication initiator protein family Orc/CDC6 identifies the primary origin of replication and serves to recruit a replicative helicase/other machinery to initiate replication.

**Archaeal-eukaryotic-like primases (AEP) primase**

Firmicutes plasmids such as pSM19035, pMD101, pBT233 encode a specialized replication protein that resembles enzymes of the superfamily of archaeo-eukaryotic primases (AEPs)[12]. Primases synthesize short RNA primers to facilitate chain elongation during replication.

**DnaB replication initiation**

Chlamydial plasmids such as the Chlamydia trachomatis plasmid encode a DnaB-like protein which is necessary for replication[13–15].

### Supplementary Note 2

The putative Methanosarcina plasmids were recovered mostly from Sphagnum acidic bogs **(Supplementary Fig. 11)**. Sphagnum acidic bogs are anaerobic environments that accumulate acetate from Sphagnum degradation, providing conditions that are conducive for acetate-utilising Methanosarcinaceae[16,17]. The associated MAG, MFD01716.bin.1.47, was predicted to support acetoclastic and hydrogenotrophic methanogenesis, but not methylotrophic methanogenesis, potentially reflecting an adaptation to these environments **(Supplementary Table 6; Supplementary Table 7; Supplementary Fig. 10)**. In all methanogenic pathways, the final methane-forming reaction converts $CH_3$-S-CoM and HS-CoB into $CH_4$ and the heterodisulfide CoM-S-S-CoB[18]. Sustained methanogenesis depends on heterodisulfide reductase (Hdr) to regenerate HS-CoM and HS-CoB by reducing CoM-S-S-CoB[18] **(Fig. 5B).**

Vht encoded by P1 could support this regeneration by oxidising $H_2$ and transferring the electrons to methanophenazine (MP) to form $MPH_2$, which HdrDE subsequently oxidises to reduce CoM-S-S-CoB (Fig. 5B). The repression of Vht in Methanosarcina barkeri prevented the recapture of $H_2$ produced during internal hydrogen cycling, resulting in $H_2$ accumulation and eventual cell lysis[19]. Accordingly, *M. barkeri* mutants lacking Vht failed to grow on $H_2/CO_2$ or acetate, whereas a mutant lacking all three active hydrogenases (Frh, Vht and Ech) retained the ability to grow on methanol, although at a reduced rate[20]. Both $H_2/CO_2$ and acetate are available to methanogens in acidic bogs[16,17]. Thus, the plasmidic Vht paralog may enhance fitness by facilitating cofactor redox cycling through gene dosage effects or through differences in activity under variable $H_2$ availability **(Fig. 5B)**. CoM-S-S-CoB can also be reduced by the cytoplasmic HdrA2B2C2 using reduced ferredoxin[21] **(Fig. 5B)**. By encoding HdrC2, the plasmid completes this complex together with the chromosomal HdrA2 and HdrB2 subunits and could thereby provide an ancillary cytoplasmic route for regenerating HS-CoM and HS-CoB **(Fig. 5B)**. HdrA2B2C2 is ubiquitous under acetotrophic growth of acetotrophic methanogens and has been proposed to become increasingly important as acetate becomes limiting[21], consistent with a modified strain of *M. acetivorans* that does not produce HdrA2B2C2 being unaffected on most substrates but growing more slowly on acetate[18,22]. Encoding HdrC2 on P2 may have increased its gene dosage, facilitating adaptation to changing electron-transfer demands compared with a single chromosomal copy. Selection could thereby favour HdrC2 variants suited to different electron-transfer demands without altering the more evolutionarily constrained catalytic HdrA2 and HdrB2 subunits. HdrC2 mediates electron transfer between these subunits, and although transfer can occur without it, the rate is substantially lower[21]. The Fe and Zn transporters could further support methanogenesis by supplying metals required by Hdr and other metalloenzymes[23]. The AglB and disaggregatase-related proteins encoded by P2 may together contribute to host ecological adaptability by facilitating transitions between aggregated and dispersed growth **(Fig. 5B)**.

The associated MAG encodes Vht, Frh and Ech but lacks Rnf, consistent with Type I Methanosarcina[24], which typically grows as aggregates embedded in a methanochondroitin matrix[25,26]. AglB is required for surface-associated growth in M. maripaludis[27], whereas disaggregatases promote dispersal as single cells in M. mazei[28]. If these proteins perform analogous functions in the associated host, the disaggregatase could release individual cells from existing aggregates, while AglB could support their attachment to new surfaces and the establishment of new aggregates. This could allow the host to colonise new environments while increasing opportunities for the plasmid to persist and spread. Why these functions are plasmid- rather than chromosome-encoded remains unclear. Selection may favour plasmids that alleviate constraints on core metabolism by increasing gene dosage, while plasmid-borne homologs could diversify without compromising the chromosome-encoded function. This flexibility could support local niche adaptation by tuning functions to prevailing conditions or promote niche expansion through specialised variants or improved acquisition of limiting metals. Although hypothetical, these mechanisms provide a rationale for encoding such functions on plasmids and suggest that the mobilome may form an adaptable extension of the *Methanosarcina* genome. This is notable in a genus whose acetoclastic metabolism was itself shaped by horizontal gene acquisition[29,30].
